# Neural hierarchy for coding articulatory dynamics in speech imagery and production

**DOI:** 10.1101/2025.05.27.656311

**Authors:** Zehao Zhao, Zhenjie Wang, Yan Liu, Youkun Qian, Yuan Yin, Xiaowei Gao, Binke Yuan, Shelley Xiuli Tong, Xing Tian, Gao Chen, Yuanning Li, Junfeng Lu, Jinsong Wu

## Abstract

Mental imagery is a hallmark of human cognition, yet the neural mechanisms underlying these internal states remain poorly understood. Speech imagery—the internal simulation of speech without overt articulation—has been proposed to partially share neural substrates with actual speech articulation. However, the precise feature encoding and spatiotemporal dynamics of this neural architecture remain controversial, constraining the understanding of mental states and the development of reliable speech imagery decoders. Here, we leveraged high-resolution electrocorticography recordings to investigate the shared and modality-specific cortical coding of articulatory kinematic trajectories (AKTs) during speech imagery and articulation. Applying a linear model, we identified robust neural dynamics in frontoparietal cortex that encoded AKTs across both modalities. Shared neural populations across the middle premotor cortex, subcentral gyrus, and postcentral-supramarginal junction exhibited consistent spatiotemporal stability during the integrative articulatory planning. In contrast, modality-specific populations for speech imagery and articulation were somatotopically interleaved along the primary sensorimotor cortex, revealing a hierarchical spatiotemporal organization distinct from shared encoding regions. We further developed a generalized neural network to decode multi-population neural dynamics. The model achieved high syllable prediction accuracy for speech imagery (79% median accuracy), closely matching the performance of speech articulation (81%). This model robustly extrapolated AKT decoding to untrained syllables within each modality while demonstrating cross-modal generalization across shared populations. These findings uncover a somato-cognitive hierarchy linking high-level supramodal planning with modality-specific neural manifestation, revolutionizing an imagery-based brain-computer interface that directly decodes thoughts for synthetic telepathy.

## Introduction

Mental imagery is a unique adaptive trait of human cognition for linking past experiences, current states, and future scenarios by simulating and creating mental events ^1-5^. For example, mental imagery of speech, also known as covert or inner speech, represents the internally generated, quasi-perceptual experience of speech without overt or audible articulation ^6,7^. This phenomenon exemplifies both an instantiation of language as a vehicle for thought and a specific form of environment-disengaged action ^8^. Existing research has shown that the neural processes of overt speech articulation involve a multi-stage sequence that comprises conceptual formulation, lexical selection, phonological encoding, motor planning, and the execution of articulatory movements ^6,7,9,10^. However, the neural mechanisms governing speech imagery are not yet fully understood. Current theories propose that speech imagery may constitute a truncated form of overt speech production, but the specific stage at which this process diverges remains debated ^6,11-13^. The abstraction view posits that speech imagery arises from the activation of abstract linguistic constructs, independent of articulatory representations ^6,11,14,15^. In contrast, the motor simulation hypothesis suggests that speech imagery mirrors overt speech articulation, involving the planning and prediction of articulatory details that are halted before execution ^6,11,13^. Clarifying this debate is critical for understanding how speech motor control interfaces with internal cognitive states.

Emerging neurolinguistic models, supported by findings from functional magnetic resonance imaging (fMRI), magnetoencephalography (MEG), and scalp electroencephalography (EEG), suggest that initial processes involved in motor planning for both speech imagery and articulation exhibit considerable similarities ^7,16-18^. Functional localization studies reveal overlapping frontoparietal activation in regions such as the premotor cortex and supramarginal gyrus (SMG), along with the articulation-specific engagement of the primary sensorimotor cortex (SMC) ^6,16-21^. Compared to actual articulation, the neural operations during speech imagery appear to suspend downstream output execution, retaining only an efference copy of motor commands ^16-18^. This copy is passed to an internal forward model in the frontoparietal cortex to facilitate internal sensorimotor simulation and predict the subsequent outcomes of speech ^16-18^. However, the precise neural hierarchy encoding internal and external articulatory movements remains unsolved, primarily due to the limited spatiotemporal resolution of non-invasive techniques and a paucity of high-precision intracranial recordings during speech imagery.

Beyond theoretical implications, speech imagery offers potential for neural decoding, particularly in restoring communication for individuals with severe speech impairments (e.g., stroke, amyotrophic lateral sclerosis [ALS], or locked-in syndrome) ^22^. Recent intracranial EEG studies have made strides in decoding vocalized, attempted, and mimed speech by leveraging articulatory movement encoding in the frontoparietal cortex ^23-31^. Yet, these approaches rely on residual motor execution signals, limiting their utility for patients with complete loss of speech output ^22,32^. Decoding speech imagery may overcome this constraint by targeting internally generated speech processes, presenting a more adaptable solution for communication restoration.

Here, we employed high-density electrocorticography (ECoG) recordings from nine participants to dissect the encoding of articulatory kinematic trajectories (AKTs) during syllable articulation (SA) and syllable imagery (SI). AKTs transcend categorical approaches by providing dynamic articulatory representations that both resolve theoretical debates (abstraction vs. motor simulation) and establish a biologically grounded foundation for naturalistic speech decoding ^28^. Using a temporal receptive field (TRF) model ^28,33,34^, we identify modality-specific AKT encoding preferences across frequency bands, characterize shared and distinct neural substrates, and map their spatiotemporal and somatotopic organization. Finally, we developed a unified decoding framework for both modalities’ syllable classification, speech synthesis, and AKT reconstruction. Notably, this framework demonstrates robust cross-modal generalization, and successfully extrapolates AKT decoding to untrained syllables. Our findings reveal a hierarchical somato-cognitive architecture underpinning speech imagery and articulation while advancing clinically viable brain-computer interfaces (BCIs) that decode speech via mental imagery in individuals with a broad spectrum of speech output impairments.

## Results

### Overview of experiments

Nine subjects (S1-S9) were recruited in this study. All subjects received awake surgeries for the treatment of brain tumors. During the awake surgery, the subjects performed the syllable articulation (SA)—syllable imagery (SI) contrast task while simultaneous audio and 256-channel high-density ECoG recordings were obtained. Among the subjects, five (S1-S5) completed Paradigm 1 (**Fig. 1a**), which involved 60 repetitions of articulation and silent imagery of six syllables, each initiated by an ipsilateral hand key press. To control for any potential influence of hand motor responses, 40 repetitions of ipsilateral key presses were performed. Electrodes responsive solely to these key presses were excluded from further analyses. The remaining four subjects (S6-S9) completed Paradigm 2 (**Fig. 1b**), which consisted of 45 repetitions of articulation and imagery tasks across eight syllables, each initiated by the appearance of a black cross symbol. To ensure the complete absence of motor and auditory output during speech imagery tasks, we implemented preoperative training combined with continuous intraoperative video/audio surveillance and electromyographic (EMG) monitoring. (Detailed in the **Methods** section)

**Fig. 1.**
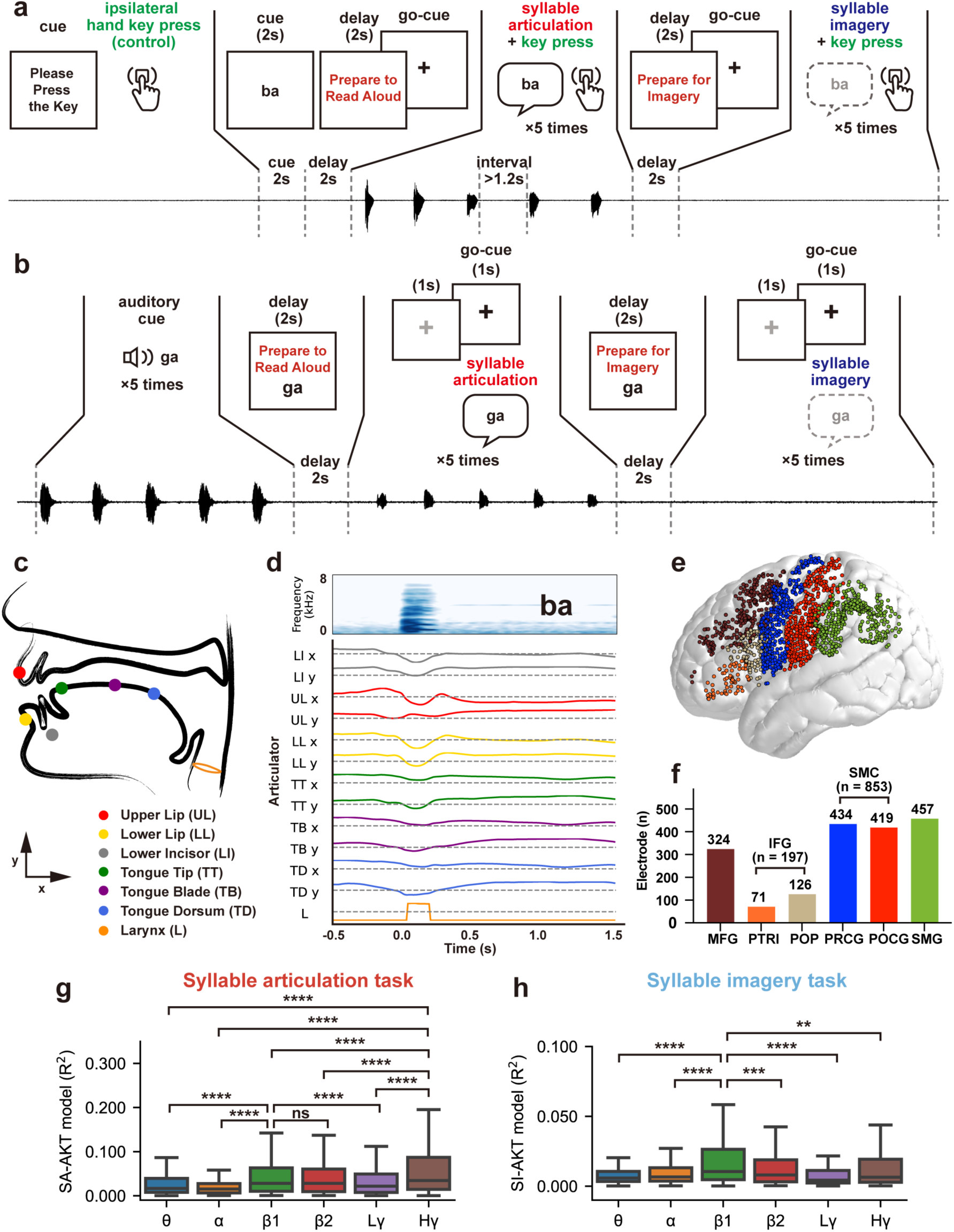
Spectral dissociation in articulatory kinematic trajectory (AKT) encoding between syllable articulation (SA) and syllable imagery (SI). **a,** Schematic of Paradigm 1. The task begins with a “Press Key” prompt, requiring subjects to press a key with the ipsilateral hand (green) 40 times as a baseline control. A visual syllable stimulus (e.g., “ba”) is then presented for 2 s, followed by a 2-s “Prepare to Read Aloud” prompt. After the appearance of the black cross symbol (go-cue), subjects articulate the syllable five times (red, SA task), marking onset with an ipsilateral key press (green, interval > 1.2 s). Next, a 2-s “Prepare for Imagery” prompt is shown, followed by the black cross symbol (go-cue), after which subjects imagine the syllable five times (blue, SI task), again marking onset with an ipsilateral key press (green, interval > 1.2 s). **b,** Schematic of Paradigm 2. Subjects first listen to an auditory syllable stimulus (e.g., “ga”) repeated five times. A 2-s “Prepare to Read Aloud” prompt and visual cue are followed by alternating grey and black crosses (1 s each). Subjects articulate the syllable upon the black cross (red, SA task), repeated five times. For imagery, a 2-s “Prepare for Imagery” prompt and visual cue precede alternating grey and black crosses, with subjects imagining the syllable upon each black cross (blue, SI task), repeated five times. **c,** Diagram of AKT movements for six articulators: upper lip (UL), lower lip (LL), lower incisor/jaw (LI), tongue tip (TT), tongue blade (TB), and tongue dorsum (TD) in 2-dimensional plane (x and y axes), along with laryngeal movement (L, log F_0_, scaled to -1 to 1 arbitrary units [a.u.], F_0_ refers to the fundamental frequency). **d,** Example Mel-spectrogram (upper) of “ba” converted into a 13-dimensional AKT time series (lower) via the acoustic-to-articulatory inversion (AAI) algorithm. **e,** The normalized total electrode coverage among cortical regions of interest on the MNI 152 brain template (subject *N* = 9, electrode *n* = 1,831): middle frontal gyrus (MFG, brown), pars triangularis (PTRI, orange), pars opercularis (POP, light yellow) of the inferior frontal gyrus (IFG), sensorimotor cortex (SMC), including precentral gyrus (PRCG, blue) and postcentral gyrus (POCG, red), and supramarginal gyrus (SMG, green). **f,** Bar graph depicting electrode counts per cortical region. **g-h,** Box plots of AKT model performance (*R*^2^) across frequency bands for SA **(g)** and SI **(h)**: theta (θ, 4-8 Hz), alpha (α, 8-12 Hz), beta1 (β1, 12-24 Hz), beta2 (β2, 24-40 Hz), low gamma (Lγ, 40-70 Hz), and high gamma (Hγ, 70-150 Hz). \*\**p* < 0.01, \*\*\**p* < 0.001, \*\*\*\**p* < 0.0001, ns: non-significant (two-sided Mann-Whitney *U* tests with Benjamini-Hochberg correction). Comparisons: SA (Hγ or β1 vs. other bands) and SI (β1 vs. other bands).

### Distinct spectral signatures govern AKT encoding across SA and SI modalities

We first analyzed the neural representation of the AKTs during the SA and SI tasks. Using an established speaker-independent acoustic-articulatory inversion (AAI) algorithm ^28^, we derived 13-dimensional AKTs from synchronized audio recordings during SA (**Fig. 1c-d**). These trajectories captured: (i) two-dimensional movements (x, y) of six articulators (upper/lower lips, jaw, tongue tip/body/dorsum); and (ii) one-dimensional laryngeal activity (scaled log F_0_)^28^. For SI tokens, we used the corresponding SA-derived AKTs as internal kinematic proxies.

Subsequently, we computed the analytic amplitude of the signals from the local field potential across seven frequency bands—theta (4-8 Hz), alpha (8-12 Hz), beta1 (12-24 Hz), beta2 (24-40 Hz), low gamma (40-70 Hz), and high gamma (70-150 Hz) ^21,25^. This analysis encompassed 1,831 electrodes distributed across our anatomically defined regions of interest (ROIs) (434 over precentral gyrus [PRCG], 419 over postcentral gyrus [POCG], 126 over pars opercularis [POP], 71 over pars triangularis [PTRI] of Broca’s area, 324 over middle frontal gyrus [MFG], and 457 over SMG) in nine participants (**Fig. 1e-f**). The beta1 band showed maximal modality responsiveness (SA: 60.0%; SI: 40.6% of electrodes), exceeding other bands (**Extended Data Fig. 1**).

To quantify how AKTs are represented in cortical activity, we modeled the relationship between time-varying AKTs and neural signals using a time-delayed ridge regression (TRF model) ^28,33,34^. This approach revealed striking modality-dependent encoding profiles: SA encoding peaked in high gamma band (median *R*^2^: 0.034, interquartile range [IQR]: 0.014–0.087; all *p* < 0.0001 vs other bands, Mann-Whitney *U* tests, Benjamini-Hochberg corrected), with secondary beta1 band representation (median *R*^2^: 0.028, IQR: 0.010–0.063; *p* < 0.0001 vs theta, alpha, and low gamma bands, *p* = 0.3784 vs beta2) (**Fig. 1g**). SI encoding was beta1-dominant (median *R*^2^: 0.011, IQR: 0.005–0.026; all *p* < 0.01 vs other bands, **Fig. 1h**). This spectral dissociation persisted across all frontoparietal regions (**Extended Data Fig. 2**), suggesting fundamental differences in how internal simulation versus motor execution engages the speech network.

### Frontoparietal beta1 activity underpins shared and specific AKT encoding in SA and SI modalities

Given our findings on the frequency-band encoding preferences, along with extensive prior research on high gamma encoding in overt speech ^19-21,25,27,28^, our analysis focuses on the beta1 band to gain insights into speech imagery. We found that 56.9% of the 1287 responsive electrodes were responsive exclusively to SA (42.2%) or to SI (14.7%) in the beta1 band, and 43.1% of all responsive electrodes responded to both modalities (**Fig. 2a**, see **Extended Data Fig. 3** for individual-level spatial distributions). Anatomical distributions differed significantly (*p* < 0.0001, *χ²* test), with dual-responsive electrodes most prevalent in PRCG (47.9%), followed by POCG (33.9%) and SMG (22.8%) (**Fig. 2a**).

**Fig. 2.**
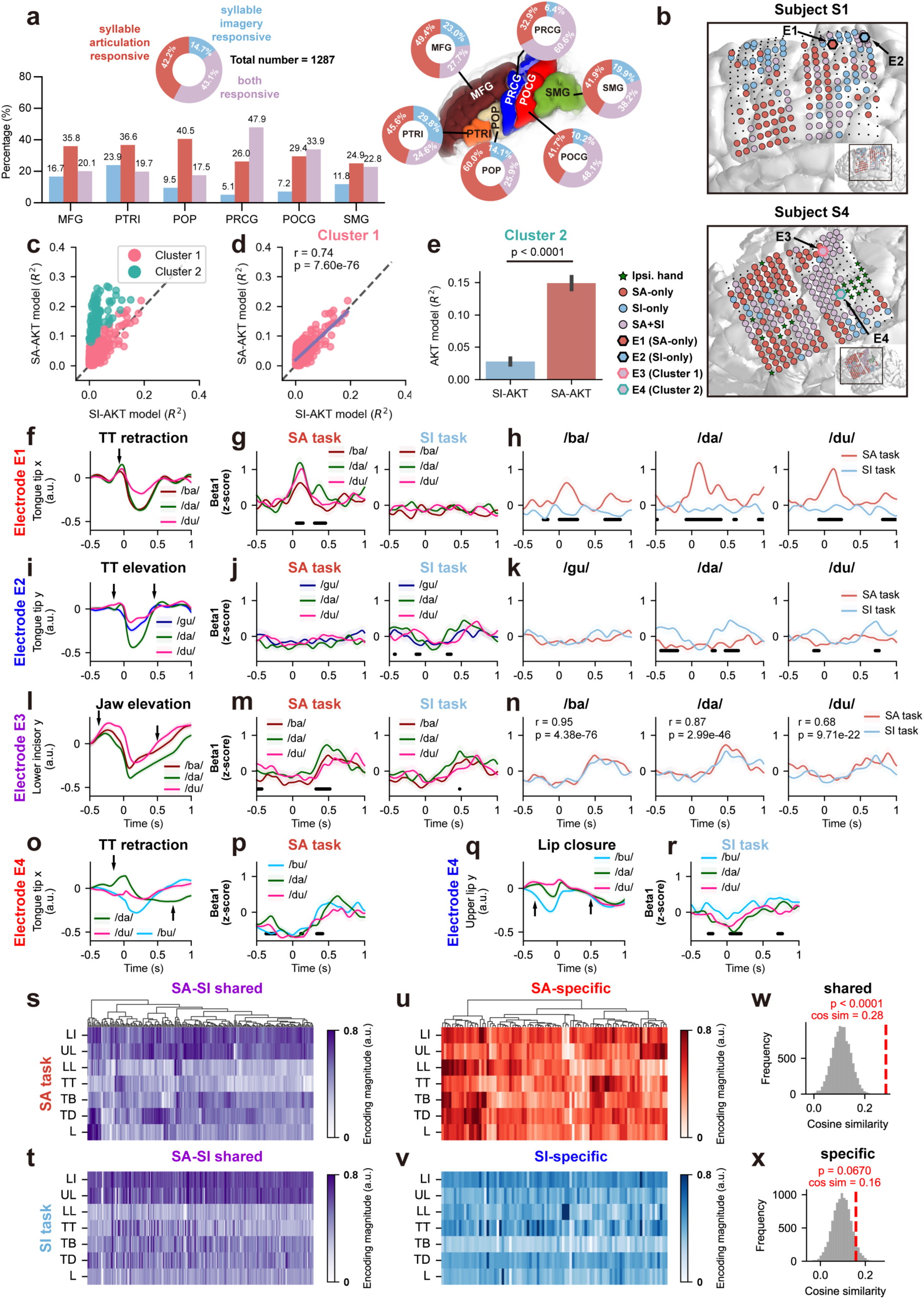
Shared and specific AKT encoding in SA and SI modalities mediated by frontoparietal beta1 activity. **a,** Electrode classification and anatomical distribution. Pie chart (top left) shows proportions of modality-selective (SA-only: red; SI-only: blue) and dual-responsive (purple) electrodes (*n* = 1,287). Bar plot (bottom left) displays the regional distribution. Right pie charts depict gyrus-specific distributions. **b,** Individual brain reconstructions for subjects S1 and S4, illustrating electrode coverage. Green stars: ipsilateral hand motor-responsive electrodes; light red, light blue, and light purple circles: SA-only, SI-only, and dual-responsive electrodes, respectively. Hexagons mark representative electrodes: E1 (SA-only, light red), E2 (SI-only, light blue), E3 (dual-responsive, Cluster 1, light purple with pink outline), and E4 (dual-responsive, Cluster 2, light purple with green outline). Black circles: non-responsive electrodes. **c,** Scatter plot illustrates the relationship between the total variance explained (*R*^2^) of the SA-AKT and SI-AKT models for each dual-responsive electrode, with colors representing the k-means clustering classification (Cluster 1: pink; Cluster 2: light green). **d,** Scatter plot highlights a strong correlation between SA- and SI-AKT model performance for Cluster 1 electrodes (*r* = 0.74, *p* = 7.60 × 10^-78^), with a purple fitted curve closely aligned to the y = x diagonal (black dashed line). **e,** Bar plot shows significant differences in SA- and SI-AKT model performance (*R*^2^, mean ± s.e.m.) for Cluster 2 electrodes (*p* < 0.0001, two-sided paired *t*-test). **f-h,** Representative SA-specific electrode (E1): tongue tip (TT) x-axis trajectories for /ba/, /da/, and /du/ (**f**) show distinct retraction magnitudes (marked by black arrow). Beta1 activity differs significantly across syllables during SA (**g**, black dots indicate time points where *p* < 0.05, one-way ANOVA) but not SI. SA beta1 activity is significantly higher than SI for all syllables (**h**, black dots indicate time points where *p* < 0.01, two-sided independent samples *t*-test). **i-k,** Representative SI-specific electrode (E2): tongue tip y-axis trajectories for /da/ and /du/ highlight unique elevations (marked by black arrows) compared to /gu/ (**i**). Beta1 activity differs significantly during SI (**j**, black dots: *p* < 0.05, one-way ANOVA) but not SA. SI beta1 activity is significantly higher than SA for /da/ and /du/ (**k**, black dots: *p* < 0.01, two-sided independent samples *t*-test), while no significant difference is observed for /gu/. **l-n,** Representative SA-SI shared encoding electrode (E3, Cluster 1): lower incisor y-axis trajectories (**l**) show varying degrees of deliberate jaw elevation (indicated by black arrows) for /ba/, /da/, and /du/. Beta1 activity differs across syllables in both tasks (**m**, black dots: *p* < 0.05, one-way ANOVA) but shows no significant differences between modalities (**n**). Instead, a strong correlation is observed between tasks (Pearson’s *r* and *p* values annotated). **o-r,** Representative Cluster 2 electrode (E4): modality-specific encoding electrode. SA-specific beta1 activity (**p**) encodes unique tongue tip retraction (**o**, marked by black arrows) for /da/ and /du/ compared to /bu/, while SI-specific beta1 activity (**r**) highlights distinct upper lip closure (**q**, marked by black arrows) for /bu/ compared to /da/ and /du/. Significant differences were marked (black dots: *p* < 0.05, one-way ANOVA). Time 0 ms indicates syllable onset (SA) or inferred onset (SI). Solid lines: mean; shaded areas: s.e.m. across syllable repetitions (color-coded). **s-t, w,** AKT encoding patterns derived from the temporal receptive field (TRF) model for SA-SI shared encoding electrodes during SA (**s**) and SI (**t**) tasks. Electrodes in (**s**) are organized by hierarchical clustering; the same order is maintained in (**t**). **(w)** Histogram of permutation testing demonstrates significant AKT encoding pattern similarity between SA-SI shared encoding electrodes during SA and SI tasks (red dashed line, *cosine similarity* = 0.28, *p* < 0.0001). **u-v, x,** AKT encoding patterns of SA-specific (**u**) and SI-specific (**v**) electrodes show no significant similarity (*p* = 0.0674, **x**) based on permutation testing (red dashed line, *cosine similarity* = 0.16, *p* = 0.0670). The color intensity in (**s-v**) reflects the unique *R*^2^ to the full TRF models (scaled to 0-1 a.u.).

Applying stringent selection criteria ( *R*^2^> 0.01, top 50% of responsive electrodes; **Extended Data Fig. 4b**), we identified functionally distinct neural populations exhibiting differential encoding of AKTs across speech modalities. SA-specific electrodes exhibited beta1 activity that precisely tracked actual articulator movements. For example, Electrode E1 (**Fig. 2b**, light red hexagon with black border) showed activity patterns (**Fig. 2g-h**) that scaled with tongue retraction demands, being strongest for alveolar /da/, intermediate for /du/, and weakest for bilabial /ba/ (**Fig. 2f**). These electrodes remained silent during imagery (**Fig. 2g-h**), revealing their exclusive role in motor execution.

Strikingly, we identified a distinct population of SI-specific encoding electrodes that maintained precise encoding of AKTs despite a complete absence of movement. Representative Electrode E2 (**Fig. 2b**, light blue hexagon with black border) exhibited significant beta1 activity (**Fig. 2j-k**) that tracked the internal dynamics of tongue tip elevation (**Fig. 2i**) during mental imagery. Specifically, the double-peaked beta1 activity pattern mirrored the kinematic sequence of simulated alveolar contact (/d/) followed by vowel-specific repositioning (**Fig. 2i-k**), demonstrating accurate internal simulation of articulation.

Our investigation of dual-responsive electrodes revealed two distinct patterns of AKT encoding across tasks. Quantitative clustering analysis (Duda-Hart statistic: *d* > 1.645, *p* < 0.05; silhouette score-optimized, **Extended Data Fig. 4a**) identified two functionally segregated populations within the *R*^2^ value distribution space (SA-AKT vs SI-AKT models, **Fig. 2c**). The first subset of electrodes (Cluster 1, pink dots in **Fig. 2c-d**) exhibited a strong positive correlation between SA-AKT and SI-AKT *R*^2^values (Pearson’s *r* = 0.74, *p* < 0.0001), suggesting shared encoding across SA and SI modalities. Representative Electrode E3 (**Fig. 2b**, light purple hexagon with pink border) maintained consistent beta1 activity for sequential jaw elevation planning of /ba/, /da/, and /du/ syllables across both modalities (*r* = 0.95 for /ba/, *r* = 0.87 for /da/, *r* = 0.68 for /du/, all *p* < 0.0001, **Fig. 2n**). These populations faithfully tracked intended articulatory demands, showing enhanced activity for open vowel /a/ compared to close vowel /u/ (**Fig. 2l-m**).

The other subset of electrodes (Cluster 2, light green dots in **Fig. 2c**), on the contrary, showed significant but distinct AKT encoding performances between SA and SI tasks (*R*^2^: 0.149 ± 0.005 vs. 0.028 ± 0.002, *p* < 0.0001, two-sided paired *t*-test, **Fig. 2e**). Representative Electrode E4 (**Fig. 2b**, light purple hexagon with light green border) switched functional specialization between modalities: during SA task it encoded tongue tip retraction (alveolar /d/ > bilabial /b/; **Fig. 2o-p**), while during SI task it preferentially represented lip closure (bilabial /b/ > alveolar /d/; **Fig. 2q-r**). This functional switching occurred with precise temporal coupling to expected articulator kinematics in each modality (**Fig. 2p, r**).

Taking a closer look at the coding properties in individual electrodes from these two clusters, we further confirmed these distinct SA-SI AKT coding patterns. For each cluster, we calculated each articulator’s unique contribution (Δ*R*^2^) to the full AKT model across electrodes. Cluster 1 encoding electrodes (e.g., Electrode E3) maintained robust cross-modal consistency between the SA modality (**Fig. 2s**) and the SI modality (**Fig. 2t**), with a *cosine similarity* of 0.28 (**Fig. 2w**, *p* < 0.0001, permutation test). This stability suggests these neural ensembles participate in a supramodal speech planning network that transcends production modality. Conversely, in Cluster 2 encoding electrodes, no significant similarity was observed between the AKT encoding patterns of the SA modality (**Fig. 2u**) and the SI modality (**Fig. 2v**) (**Fig. 2x**, *cosine similarity* = 0.16, *p* = 0.0674, permutation test). This modality-specific specialization likely reflects microanatomical segregation of functionally distinct neural subpopulations within the same cortical territory, with differential engagement based on behavioral state (e.g., Electrode E4, serving as SA-specific electrodes in SA modality and as SI-specific electrodes in SI modality) ^35^.

Our results demonstrate that speech imagery constitutes an active cognitive process involving detailed internal articulatory representations, distinct from truncated articulation. While partial overlap with speech articulation (Cluster 1, Electrode E3) confirms shared planning mechanisms ^6,7,9,10^, the discovery of modality-specific neural signatures—particularly SI-exclusive AKT encoding (Electrode E2) and Cluster 2’s functional switching (Electrode E4)—reveals unique computational principles governing imagined speech. Identifying these internal kinematic codes provides a neurophysiological basis for decoding imagined speech, with immediate implications for brain-computer interfaces to restore communication in paralysis.

### Spatiotemporal hierarchy separates supramodal and modality-specific articulatory representations

To investigate the spatial distribution patterns of SA-SI shared, SA-specific, and SI-specific encoding electrodes at the population level, we normalized all encoding electrodes to the MNI 152 template and applied kernel density estimation (KDE) to construct probability density distributions for each encoding category (**Fig. 3a–c**). SA-specific encoding electrodes (**Fig. 3a, e**) and SI-specific encoding electrodes (**Fig. 3c, h**) were concentrated bilaterally around the central sulcus, predominantly covering the ventral primary sensorimotor cortex (areas 4, 3b, 1, and 2, according to a multi-modal parcellation atlas ^36^). The peak point for SA-specific electrodes was located posteroinferior to that of SI-specific electrodes (**Fig. 3m**).

**Fig. 3.**
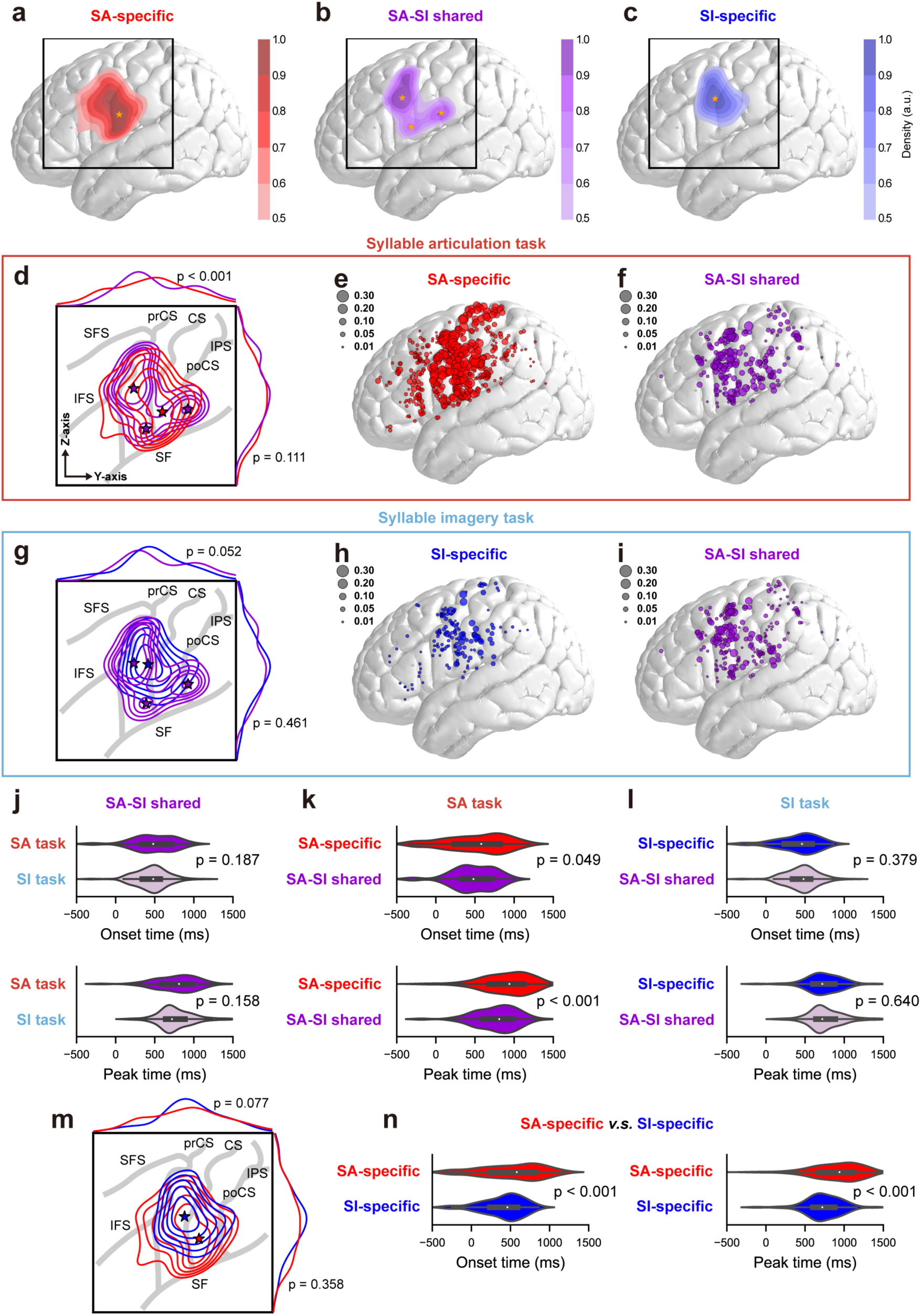
Spatial distribution and temporal dynamics of SA-specific, SI-specific, and SA-SI shared encoding electrodes. **a-c,** Probability density maps of SA-specific encoding electrodes (red, **a**), SA-SI shared encoding electrodes (purple, **b**), and SI-specific encoding electrodes (blue, **c**) using kernel density estimation (KDE). Darker colors indicate higher probability density, with a cut-off at 0.5 cumulative density (a.u.). Yellow stars mark the peak density points, and black boxes indicate the regions shown in panels **d**, **g**, and **m**. **d,** KDE contour plot comparing SA-specific (red) and SA-SI shared (purple) encoding electrodes during the SA task. KDE curves along the Y- and Z-axes of MNI space are shown, with *p*-values from two-sided Kolmogorov-Smirnov tests. The gray solid lines mark key brain sulci: prCS = precentral sulcus, CS = central sulcus, poCS = postcentral sulcus, SF = Sylvian fissure, SFS = superior frontal sulcus, IFS = inferior frontal sulcus, IPS = intraparietal sulcus. **e-f,** Spatial scatter plots of SA-specific encoding electrodes (**e**, red) and SA-SI shared encoding electrodes (**f**, purple) during the SA task. The size of each point reflects the *R*^2^ value in the SA-AKT model. **(g)** KDE contour plot comparing SI-specific (blue) and SA-SI shared (purple) encoding electrodes during the SI task. KDE curves are displayed along the Y- and Z-axes with *p*-values from two-sided Kolmogorov-Smirnov tests. **h-i,** Spatial scatter plots of SI-specific encoding electrodes (**h**, blue) and SA-SI shared encoding electrodes (**i**, purple) during the SI task. Point size reflects the *R*^2^value in the SI-AKT model. **j,** Violin plots illustrating the onset (top) and peak (bottom) times of SA-SI shared encoding electrodes during the SA task (purple) and SI task (light purple), showing no significant differences based on two-sided Mann-Whitney *U* tests. **k,** Violin plots showing the onset (top) and peak (bottom) times of SA-specific (red) and SA-SI shared (purple) encoding electrodes during the SA task, with later timings for SA-specific electrodes based on two-sided Mann-Whitney *U* tests. **l,** Violin plots showing the onset (top) and peak (bottom) times of SI-specific (blue) and SA-SI shared (purple) encoding electrodes during the SI task, with no significant differences observed based on two-sided Mann-Whitney *U* tests. **m,** KDE contour plot comparing the spatial distribution of SA-specific (red) and SI-specific (blue) encoding electrodes. KDE curves are displayed along the Y- and Z-axes with *p*-values from two-sided Kolmogorov-Smirnov tests. **n,** Violin plots comparing the onset (left) and peak times (right) of SA-specific (red) and SI-specific (blue) encoding electrodes during their respective tasks, with later timings observed for SA-specific electrodes based on two-sided Mann-Whitney *U* tests.

In contrast, supramodal (SA-SI shared encoding) electrodes (**Fig. 3b**) exhibited three distributed clusters (**Fig. 3b**): (1) middle premotor cortex (spanning areas 55b and dorsal 6v with peak density in dorsal 6v), (2) subcentral gyrus (encompassing areas 43 and ventral 6v centered on area 43), and (3) supramarginal-postcentral junction (covering areas OP4, PFop, and PF with focal concentration in OP4). These three peaks of supramodal electrodes were positioned anterosuperior, anteroinferior, and posteroinferior relative to those of SA-specific and SI-specific electrodes (**Fig. 3d, g**), exhibiting y-axis displacement (two-sided Kolmogorov-Smirnov test: *p* < 0.001 vs SA-specific; *p* = 0.052 vs SI-specific) (**Fig. 3d–i**).

Next, we analyzed the temporal dynamics of the different electrode groups, specifically comparing across electrode groups for the onset and peak times in SA and SI tasks. Our analysis revealed distinct temporal activation patterns across neural populations (**Fig. 3j-n**). Supramodal (SA-SI shared encoding) electrodes showed remarkable temporal stability, with no significant difference in median onset time (median [IQR], SA: 480 ms [328–710 ms], SI: 480 ms [340–570 ms], *p* = 0.187, two-sided Mann-Whitney *U* test) or peak time (SA: 810 ms [590–980 ms], SI: 720 ms [640–885 ms], *p* = 0.158) between modalities (**Fig. 3j**). During SA task, these supramodal electrodes activated significantly earlier than SA-specific electrodes (onset: 580 ms [235–820 ms], 100 ms difference, *p* = 0.049; peak: 940 ms [680–1120 ms], 130 ms difference, *p* < 0.001), forming a temporally cascaded processing stream (**Fig. 3k**). During the SI task, SI-specific electrodes showed synchronous activation (onset: 460 ms [235–590 ms], *p* = 0.379; peak: 720 ms [600–880 ms], *p* = 0.640; **Fig. 3l**) with supramodal electrodes. In contrast, SA-specific electrodes exhibited consistently delayed activation compared to SI-specific electrodes across all temporal measures (all *p* < 0.001), revealing fundamentally different processing architectures between production modalities (**Fig. 3n**).

These findings suggest a hierarchical processing model where supramodal populations in middle premotor cortex, subcentral gyrus, and supramarginal-postcentral junction represent a common upstream for abstract articulatory planning. During overt speech articulation, this supramodal activity cascades to SA-specific populations in the ventral primary sensorimotor cortex for motor execution. During mental imagery, concurrent SI-specific activation across the same regions preserves internal articulatory output, enabling purely cognitive speech simulation while generating quasi-perceptual experience ^6^.

### Supramodal integration versus modality-specific somatotopic organization of articulatory gestures

Previous ECoG studies have revealed that neural dynamics at the single-electrode level encode a diverse repertoire of AKTs during continuous speech production ^28^, each reflecting coordinated movements of articulators toward four primary vocal tract shapes corresponding to distinct places of articulation (POAs): coronal, labial, dorsal constrictions, and vocalic control. Based on this, we hypothesized that beta1 activity in SA and SI modalities exhibits similar encoding modes.

To test this hypothesis, we first examined whether AKTs associated with different phonemes could be organized according to the similarity of their articulatory gestures. We computed the displacement matrix of AKTs during articulation and performed unsupervised hierarchical clustering across phonemes. This analysis revealed a clear grouping of phonemes into four established POA categories, with subtle intra-cluster variations (**Fig. 4a**). We then aggregated AKTs within each POA-defined cluster and found that core articulatory features were preserved, reinforcing the consistency of gesture patterns within each category (**Fig. 4d**).

**Fig. 4.**
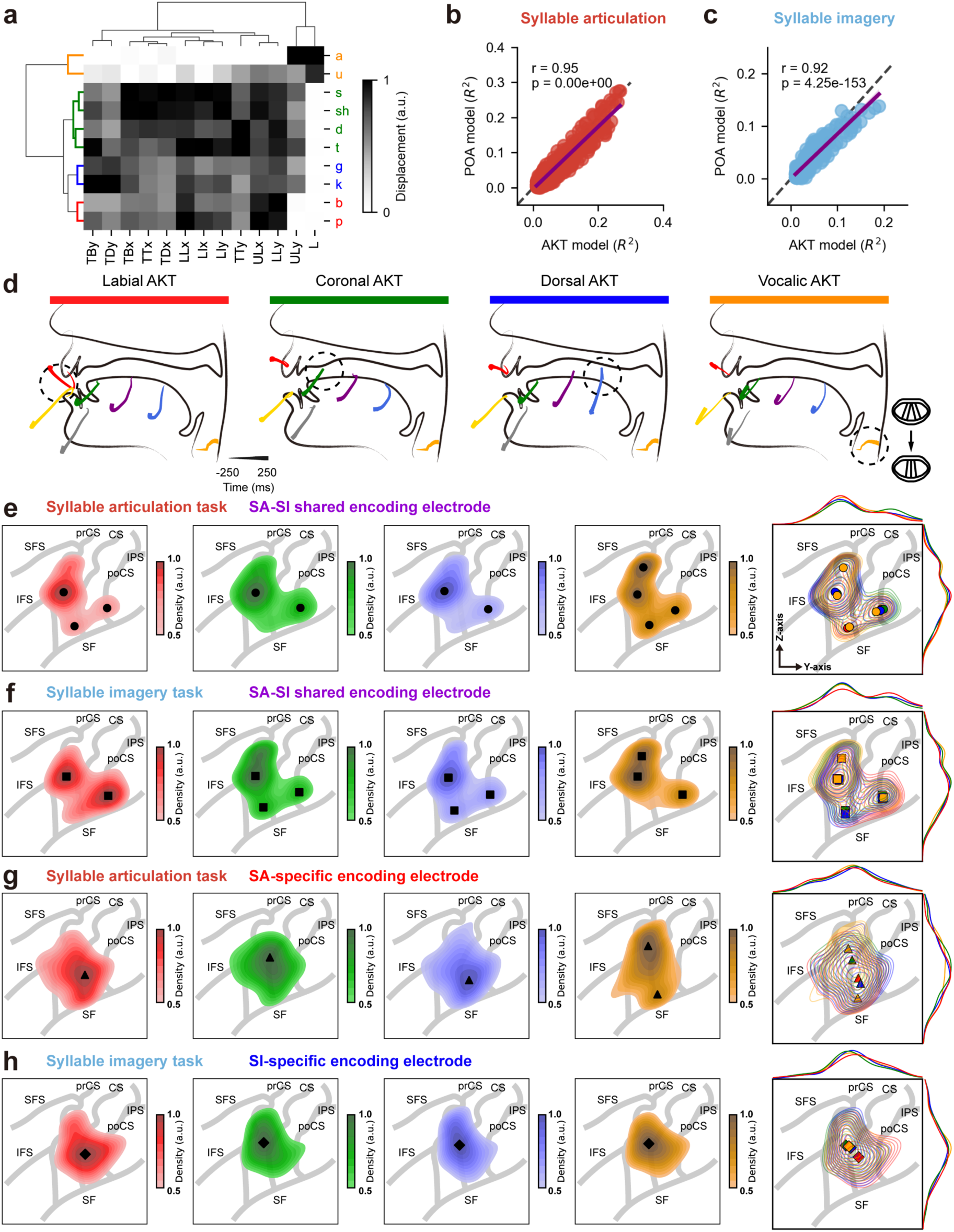
Somatotopic arrangement of four distinct vocal tract gestures in SA-specific, SI-specific, and SA-SI shared encoding electrodes. **a,** Hierarchical clustering of phonemes based on AKT displacement follows the place of articulation (POA) categories (red: labial, green: coronal, blue: dorsal, orange: vocalic). **b-c,** Scatter plots show strong correlations between the total variance explained (*R*^2^) of the 13-dimensional AKT and 4-dimensional POA-based TRF models during both syllable articulation (**b**) and imagery (**c**) tasks (SA: Pearson’s *r* = 0.95, *p* < 0.0001; SI: Pearson’s *r* = 0.92, *p* < 0.0001), with purple fitted curves closely aligned to the y = x diagonal (black dashed line). **d,** Averaged movement trajectories of seven articulators grouped by four categories of POA during phoneme articulation across all nine subjects. The time window spans from -250 ms to 250 ms relative to phoneme onset. The time course of trajectories is represented by thin-to-thick lines. Dashed circles highlight the primary vocal tract constriction for each POA category: labial AKT (red) involves lip closure, coronal AKT (green) involves tongue tip elevation contacting the alveolar ridge, dorsal AKT (blue) involves tongue dorsum elevation against the soft palate, and vocalic AKT (orange) involves the constriction and approximation of the vocal folds for voicing. **e-h,** Weighted KDE probability density distributions for the four POA categories during the syllable articulation task (**e, g**) and the syllable imagery task (**f, h**), visualized for SA-SI shared (**e, f**), SA-specific (**g**), and SI-specific (**h**) encoding electrodes. In the first four columns of panels **e-h**, the distributions were weighted by the Δ*R*^2^ values (normalized to a scale of 0–1) of the POAs, with a cut-off at 0.5 cumulative density (a.u.). Each color represents a distinct POA: red for labial, green for coronal, blue for dorsal, and yellow for vocalic. The intensity of the color corresponds to the probability density, with darker shades indicating higher density. Circles (**e**), squares (**f**), triangles (**g**), or diamonds (**h**) indicate the positions of the probability density peaks. The last column of panels **e-h** displays joint weighted KDE contour plots, with colored symbols marking the peak locations for each POA. Gray solid lines indicate major brain sulci: prCS = precentral sulcus, CS = central sulcus, poCS = postcentral sulcus, SF = Sylvian fissure, SFS = superior frontal sulcus, IFS = inferior frontal sulcus, and IPS = intraparietal sulcus. Refer to **Extended Data Figs. 6 and 7** for original scatter plots, all peak locations, and pairwise permutation test results of POA encoding across SA-specific, SI-specific, and SA-SI shared encoding electrodes during SA and SI modalities.

To further evaluate the encoding of these POA features in beta1 activity, we trained a TRF model incorporating POA cluster-based features, marking the presence or absence of each phoneme’s POA feature at its onset using binary indicators (1/0). We found a strong positive correlation between the total variance explained (*R*^2^) of the 13-dimensional AKT-based TRF model and the 4-dimensional POA-based TRF model in both SA and SI modalities (SA: Pearson’s *r* = 0.95, *p* < 0.0001; SI: *r* = 0.92, *p* < 0.0001) (**Fig. 4b-c**), indicating a high degree of consistency in the neural encoding of AKT and POA representations across both modalities. This also suggests that the frontoparietal regions exhibit a relatively rigid encoding of AKT combinations, which closely aligns with the four distinct articulatory gestures.

Subsequently, we aimed to determine the somatotopic arrangements across POAs for SA-SI shared, SA-specific, and SI-specific encoding electrodes, respectively. To this end, we constructed weighted functional maps for each POA in the y-z plane of MNI space, using each POA’s unique contribution (Δ*R*^2^) as the weighting factor in the KDE to visualize the spatial encoding patterns for the three categories of electrodes.

Our spatial analysis revealed fundamentally distinct organizational principles between supramodal and modality-specific articulatory representations. For SA-SI shared encoding electrodes, we observed overlapping functional peaks for multiple POAs in three key regions: the middle premotor cortex, subcentral gyrus, and POCG-SMG junction (**Fig. 4e-f**, consistent with peak locations identified in **Fig. 3b**). This conserved spatial patterning across both speech modalities (*cosine similarity* = 0.56, *p* < 0.0001, **Extended Data Fig. 5**) suggests that these supramodal regions implement integrative planning and coordination of multiple articulatory gestures ^10,37-39^. In addition, we identified a functional peak for vocalic constriction at the MFG-PRCG junction under both SA and SI modalities, suggesting a preferential encoding for vocalic control planning in this region.

In contrast, modality-specific electrodes exhibited distinct somatotopic arrangements. SA-specific electrodes exhibited a well-defined ventral-dorsal somatotopic gradient in the vSMC, with dual vocalic representations at the ventral and dorsal extremes. Between these vocalic peaks, we observed sequential representations of coronal, labial, and dorsal POAs, with significant separation between all adjacent peaks (**Fig. 4g**; pairwise permutation test results shown in **Extended Data Fig. 6**). Furthermore, SI-specific encoding electrodes displayed a less distinct somatotopic arrangement along the central sulcus, oriented from superior-anterior to inferior-posterior, with coronal, vocalic, dorsal, and labial peaks positioned sequentially (**Fig. 4h, Extended Data Fig. 6**). However, only the separation between labial and coronal peaks reached statistical significance (*p* = 0.0336 along the MNI y-axis; pairwise permutation tests, Benjamini-Hochberg corrected, **Extended Data Fig. 6**).

Our findings reveal a hierarchical somato-cognitive architecture for speech imagery and articulation. Within distributed frontoparietal nodes, spatially condensed supramodal assemblies support compressed articulatory gesture planning, enabling computational efficiency. These abstract representations dynamically interface with modality-specific somatotopic implementations in primary sensorimotor cortex: the system maintains clear somatotopic organization during overt articulation, whereas covert imagery engages a reconfigured functional topography that facilitates internal simulation.

### Frontoparietal articulatory codes enable effective syllable classification, speech synthesis, and kinematic decoding across executed and imagined speech

Having characterized the encoding patterns of AKTs and POAs at the single-electrode level for both modalities, we next asked whether population-level articulatory codes in frontoparietal cortex could support speech decoding, particularly for inner speech in the SI condition. To address this, we developed a triple-stream neural decoding framework (**Fig. 5a**) that maps beta1 activity from SA- and SI-responsive electrodes to three parallel outputs: syllable classification (syllable classifier; **Fig. 5b**), acoustic waveform reconstruction (speech synthesizer; **Fig. 5c**), and AKTs (articulatory movement synthesizer; **Fig. 5d**). Decoder performance was evaluated across all nine subjects using syllable classification accuracy, Mel cepstral distortion (MCD) for synthesized audio, and Pearson’s correlation coefficient (*r*) for predicted articulatory trajectories.

**Fig. 5.**
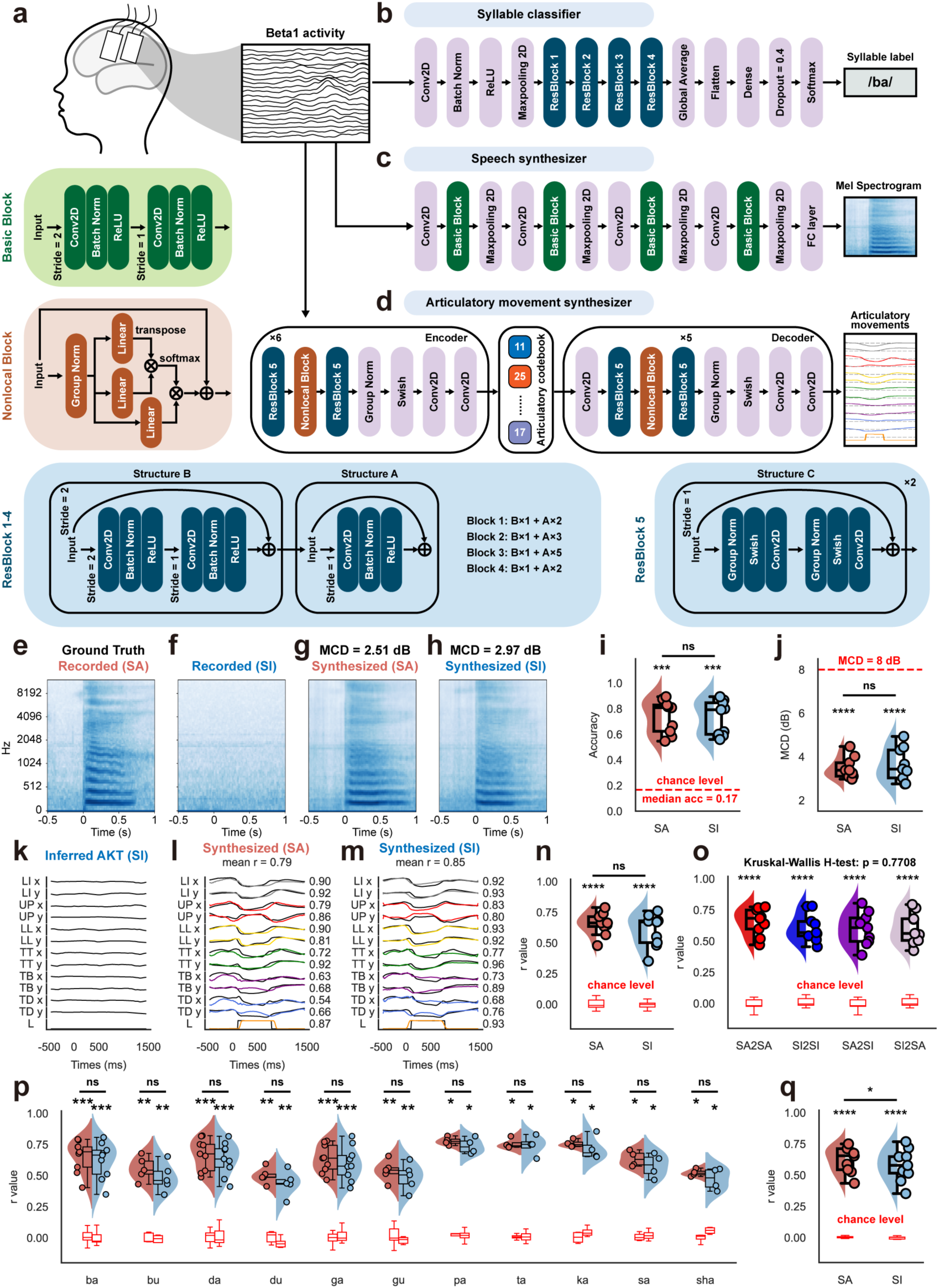
Deep learning-based synthesis of text, speech, and articulatory kinematics from frontoparietal beta1 activity during SA and SI tasks. **a,** Schematic diagram of the deep learning model pipeline for synthesizing text, speech, and AKTs from frontoparietal beta1 activity. **b-d,** Model architectures for the syllable classifier (**b**), speech audio synthesizer (**c**), and articulatory movement synthesizer (**d**). **e,** Recorded Mel-spectrogram of a /ba/ during the SA task (ground truth). **f,** Recorded Mel-spectrogram during syllable imagery of a /ba/ (blank control). **g-h,** Mel-spectrogram synthesized from beta1 activity recorded during SA (**g**), and SI (**h**) modalities, with Mel-Cepstral Distortion (MCD) values of 2.51 dB and 2.97 dB, respectively. **i,** Violin plots of syllable classifier accuracy for electrodes responsive to SA and SI tasks (subject *N* = 9). The red dashed line indicates the median chance level; \*\*\**p* < 0.001 for Mann–Whitney *U* test against chance level; ns: non-significant, Wilcoxon signed-rank test for SA vs. SI. **j,** Violin plot of synthesized speech MCD values. The red dashed line represents the 8 dB threshold considered acceptable for voice recognition; \*\*\*\**p* < 0.0001 for Mann–Whitney *U* test against the threshold; ns: non-significant, Wilcoxon signed-rank test for SA vs. SI. **k,** AKTs inferred from audio recorded during SI modality using AAI algorithm (blank control). **l-m,** Synthesized AKTs from beta1 activity following Leave-One-Syllable-Out strategy in SA (**l**) and SI (**m**) tasks, with mean Pearson’s *r* values across all AKTs of 0.79 and 0.85, respectively. Black lines indicate ground truth AKTs inferred from audio recorded during SA modality. **n,** Violin plot comparing the performance of synthesized AKTs, represented by mean Pearson’s *r* values of all AKTs with ground truth, for SA and SI-responsive electrodes (subject *N* = 9). The red box represents chance level; \*\*\*\**p* < 0.0001 for Mann–Whitney *U* test against chance level; ns: non-significant, Wilcoxon signed-rank test for SA vs. SI. **o,** Violin plot of cross-modal transfer learning performance for SA-SI shared electrodes (subject *N* = 9), with the red box marking chance level. X-axis labels are formatted as "training set to test set" (e.g., SA2SI indicates a model trained on SA and tested on SI); \*\*\*\**p* < 0.0001 for Mann–Whitney *U* test against chance level. Kruskal-Wallis H-test across models: *p* = 0.7708. **p,** Leave-One-Syllable-Out performance of all syllables for SA and SI responsive electrodes. The x-axis represents the syllables left out. The red box represents chance level; *, **, ***, **** denote Mann–Whitney *U* test significance at *p* < 0.05, 0.01, 0.001, and 0.0001, respectively, against chance level; ns: non-significant, Wilcoxon signed-rank test for SA vs. SI. **q,** Summary violin plot for Leave-One-Syllable-Out data comparing SA and SI modalities (subject *N* = 9); \*\*\*\**p* < 0.0001 for Mann–Whitney *U* test against chance level, \**p* < 0.05 (Wilcoxon signed-rank test for SA vs. SI). All data in the violin and box plots represent median and interquartile ranges across subjects.

The syllable classifier achieved high decoding accuracy in both SA and SI tasks, with comparable performance (SI: median accuracy = 0.79, IQR = 0.60–0.85; SA: 0.81, IQR = 0.62–0.83; *p* = 0.3594, Wilcoxon signed-rank test). Crucially, decoding in the SI modality was significantly above chance (median chance = 0.17, *p* = 0.0003, Mann-Whitney *U* test), indicating that internal articulatory representations are sufficiently stable to support categorical decoding (**Fig. 5i**).

Given that no audio is generated in the SI modality (as shown in **Fig. 5f** for covertly reading /ba/), we used the audio recorded in the SA modality for the same trial within the same block as the shared ground truth (see **Fig. 5e** for overtly reading /ba/) for both the SA and SI modalities to train the speech synthesizer. The Mel cepstral distortion (MCD) between the synthesized and real speech spectrograms was significantly below the 8 dB threshold in both the SA (median MCD = 3.38 dB, IQR: 3.11–3.72 dB) and SI (median MCD = 3.43 dB, IQR: 3.03–4.31 dB) modalities (both *p* = 0.0039), which is considered acceptable for voice recognition systems ^30^. SI decoding achieved speech synthesis quality comparable to SA (*p* = 0.3008), supporting the viability of mental imagery for high-fidelity output (**Fig. 5j**; see **Fig. 5g-h** for synthesized /ba/ Mel spectrograms for both modalities).

Similarly, using inferred articulatory trajectories from SA trials as the ground truth, we decoded AKTs from beta1 activity using all SI-responsive electrodes (**Fig. 5k–m**). Synthesized trajectories in the SI modality showed strong correlations with ground truth (median *r* = 0.67, IQR = 0.50–0.68), comparable to those observed in SA (0.66, IQR = 0.63–0.71; *p* = 0.0547), and both well above chance (*p* = 0.0004) (**Fig. 5n**). Similar decoding performance was observed when using only SI-specific electrodes (**Extended Data Fig. 8**), further supporting that internal articulatory dynamics are robustly encoded in SI-related beta1 activity.

As a result, deep neural network models trained and tested within each task demonstrated equivalent decoding efficacy for SA and SI tasks. Next, we investigated whether cross-modal articulatory movement decoding could be achieved using SA-SI shared electrodes. We trained four articulatory movement synthesizers based on these electrodes: SA2SA, SA2SI, SI2SI, and SI2SA (naming format indicates "training set to testing set," e.g., SA2SI represents training with SA modality beta1 activity and testing with SI modality beta1 activity, evaluating the correlation between synthesized movement and ground truth). Overall, we found no significant difference in the median correlation coefficients (SA2SA: 0.68 [0.60–0.75], SA2SI: 0.61 [0.50–0.69], SI2SI: 0.57 [0.54–0.66], SI2SA: 0.56 [0.51–0.68], *p* = 0.7708, Kruskal-Wallis *H*-test), with all models significantly above chance level (all *p* = 0.0004, Mann-Whitney *U* test) (**Fig. 5o**). These results further support that SA-SI shared electrodes encode supramodal articulatory representations, offering a principled basis and implantation target for cross-modal speech BCIs. Such systems may be especially beneficial for individuals with progressively declining speech function, such as those diagnosed with ALS.

We next aimed to validate whether our articulatory movement synthesizer had effectively learned the stable relationship between beta1 activity and AKTs. If this were the case, the synthesizer should exhibit extrapolative generalization, meaning it should be capable of predicting the movement trajectories of syllables not included in the training set. To assess this, we employed a Leave-One-Syllable-Out strategy, iteratively excluding one syllable, training the synthesizer on the remaining syllables, and testing the excluded syllable to compute the correlation (*r*) between predicted AKTs and ground truth. We found that the Leave-One-Syllable-Out articulatory movement synthesizer still performed well for all syllables in both SA and SI responsive electrodes, with the median correlation coefficient for all syllables significantly exceeding chance levels (*p* < 0.05 for all syllables, Mann-Whitney *U* test) (**Fig. 5p**; see **Fig. 5l-m** for synthesized articulatory movements of /ba/ in SA and SI modalities using Leave-One-Syllable-Out strategy). In summary, the Leave-One-Syllable-Out model demonstrated strong performance in both the SA (median *r* = 0.65, IQR: 0.54–0.71) and SI (median *r* = 0.58, IQR: 0.52–0.65) modalities (all *p* = 0.0004 vs chance level, **Fig. 5q**). Collectively, our articulatory movement synthesizer demonstrated robust extrapolative generalization across both modalities, indicating that it successfully captured a neurocomputationally interpretable mapping between cortical signatures and intended articulatory dynamics.

## Discussion

Our high-density ECoG recordings reveal a hierarchical cortical architecture underpinning shared and modality-specific mechanisms for articulatory control in speech imagery and articulation. Three key advances emerge: First, frontoparietal AKT encoding exhibits a spectral dissociation, with high-gamma dominance and secondary beta1 contributions in SA, contrasting with a primary beta1 signature in SI. Second, beta1-band analysis identifies spatiotemporally stable supramodal populations in the middle premotor cortex, subcentral region, and postcentral–supramarginal areas that support integrated multi-articulatory planning. These populations hierarchically interface with somatotopically interleaved primary sensorimotor populations to segregate internal simulation during speech imagery from external execution during articulation. Third, neural decoding based on these neural dynamics enables robust within-modal classification, speech synthesis, and kinematic prediction, while also demonstrating cross-modal generalization and extrapolation to untrained syllables in both modalities.

### Distinct spectral signatures across speech modalities

Our findings reveal a fundamental dissociation in spectral encoding strategies between speech imagery and articulation. High-gamma activity emerges as the dominant carrier of AKT information during overt speech articulation, particularly in frontoparietal regions, consistent with its established role in encoding fine-grained articulatory features, including AKT sequences, place of articulation, and phonetic details ^19-21,25,27,28,40,41^. Notably, while residual high-gamma activity persists during speech imagery ^19-21,40^, its functional engagement is markedly attenuated—both in spatial extent (reduced electrode recruitment) and decoding performance (*R*^2^ ranked fourth among frequency bands).

Conversely, beta1 activity exhibits maximal responsiveness and reliably encodes AKTs across both speech modalities. This band not only achieves secondary decoding performance during articulation (surpassed only by high gamma) but also emerges as the principal information carrier during imagery throughout frontoparietal regions. This dual-domain involvement aligns with converging evidence from human and primate studies that implicate beta1 activity in motor planning and execution, action simulation and imagery, and postural maintenance in humans and non-human primates ^40,42-47^.

### An expanded somato-cognitive architecture for speech imagery and articulation

Recent work has reconceptualized the sensorimotor cortex as a somato-cognitive action network (SCAN), comprising interdigitated effector-specific zones and integrative inter-effector regions ^39,48^. By focusing on beta1 activity, our findings extend this ‘new homunculus’ framework by revealing functional distinctions between supramodal and modality-specific populations involved in speech planning, internal simulation and external execution.

In addition to previously described inter-effector middle premotor and subcentral hubs ^39,48^, we identify a key parietal node at the POCG–SMG junction that, together with frontal counterparts, forms an extended fronto-parietal network for articulatory planning. Our data reveals that supramodal populations within these hubs exhibit three key features: robust cross-modal stability across spatiotemporal profiles and AKT encoding patterns; spatially overlapping encoding of multiple articulatory gestures, supporting integrative control; and consistent temporal precedence over downstream somatotopically organized, modality-specific populations—by ∼100 ms at onset and ∼130 ms at peak activity. These functional signatures are aligned with direct cortical stimulation studies, which show that disruption of these peri-Sylvian regions reliably induces complete speech arrest, implicating their role in coordinated articulatory planning rather than isolated motor output ^38,49^.

Further converging evidence comes from fMRI studies showing strong connectivity between these hubs and the cingulo-opercular network (CON), a domain-general system implicated in goal-directed action and performance monitoring ^39,50,51^, as well as from anatomical studies showing their tight coupling via the superior longitudinal fasciculus-III / arcuate fasciculus complex—critical tracts within the dorsal phonological stream ^9,36,49,52-54^. Stimulation of these fiber tracts likewise disrupts speech, resulting in phonemic paraphasia or complete speech arrest ^52,55-57^. Utah array recordings have further shown that limited coverage within these hubs suffices for decoding natural speech involving complex multi-articulator coordination ^26,58,59^. Together, these findings support the interpretation that supramodal populations encode cognitive-level articulatory plans generalizable across modalities. This provides direct neurophysiological support for the motor simulation hypothesis over the abstraction view, suggesting that speech imagery entails precise, temporally structured articulatory planning that is initiated but not executed ^6,11,13^.

In contrast, modality-specific populations show distinct anatomical and functional profiles. Predominantly localized to the primary sensorimotor cortex along the central sulcus, SA- and SI-specific populations are interleaved in a mosaic-like distribution that, while spatially overlapping, diverges in temporal dynamics and cross-modal encoding patterns.

Consistent with prior findings, SA-specific encoding populations exhibited a characteristic somatotopic organization, in sharp contrast to the integrative arrangement of supramodal regions. Vocalic AKTs evoked two spatial peaks at the dorsal and ventral extremes of vSMC, with coronal, labial, and dorsal peaks arranged sequentially along the superior–inferior axis. These vocalic peaks align closely with laryngeal motor areas identified in previous ECoG and fMRI work ^27,60^. Cortical stimulation at these vocalic sites induces involuntary vocalizations, pitch change, laryngeal constriction, or paralysis, providing causal evidence for their functional specificity ^29,48,61-63^. Temporally, the activation of supramodal populations consistently preceded that of SA-specific populations, echoing the ∼145 ms gap between phonetic encoding and articulatory output observed in large-scale neuroimaging meta-analyses ^64-66^.

In contrast, SI-specific electrodes exhibited a remapped functional organization with attenuated somatotopic segregation. This reduced somatotopic differentiation likely stems from their sparsely distributed and spatially dispersed organization. This may also explain the limited detection of SI-specific activity in primary sensorimotor cortex in prior fMRI studies ^67,68^. The dominant motor signal during articulation likely masks more subtle SI-related activations in cross-modal contrasts.

Remarkably, our high-density ECoG recordings identified distinct populations that exclusively encode internal AKTs during speech imagery (e.g., **Fig. 2i–k**), providing direct neurophysiological evidence for cognitive-level articulatory outputs. These internal simulations paralleled the temporal profiles of supramodal populations and may contribute to the quasi-perceptual experience of speech ^17,18^. At the same time, they may suppress overt execution via local inhibitory interactions with neighboring SA-specific circuits, enabling mental speech simulation without triggering actual articulation ^7,10,17,18,69-71^.

Together, these findings extend the SCAN framework to a broader fronto-parietal architecture linking overt articulation and covert imagery. Within this structure, higher-level, supramodal articulatory planning regions interface with modality-specific execution systems, forming a hierarchical somato-cognitive map in the sensorimotor cortex. This hierarchy is mirrored by anatomical gradients in myelin content, receptor density, and cytoarchitectonic differentiation, underscoring the specialized contributions of these regions to speech control across internal and external domains ^36,72-74^.

### Implications for neurally interpretable within- and cross-modal speech BCIs

Our encoding results establish the basis for a biologically grounded framework for decoding both spoken and imagined speech. Despite the constraints of a limited ECoG dataset, we developed a triple-stream architecture that accurately classified syllables, synthesized acoustic output, and predicted AKTs by leveraging frontoparietal beta1 activity—an underutilized neural signal in prior BCI efforts. Within each modality, this framework achieved performance levels comparable to those reported in intraoperative and early-stage implanted BCI studies ^20,21,23,58,59,75^, with considerable potential for further improvement ^25,26,76^, particularly in decoding speech imagery.

A key advance underlying this framework is the discovery of supramodal articulatory representations within the frontoparietal cortex. These representations exhibited robust cross-modal encoding stability, enabling generalization between modalities with prediction consistency approaching within-modality levels. This degree of cross-modal transferability suggests a shared speech planning architecture, capable of integrating both overt articulation and covert speech imagery. Such systems may be particularly valuable for individuals with progressive speech impairment, including those affected by ALS. Critically, the localization of these supramodal representations to the middle premotor cortex, subcentral gyrus, and POCG–SMG junction provides anatomically and functionally grounded targets for future cross-modal BCI implantation. More broadly, by engaging higher-order speech planning nodes (e.g., the SMG node) and bypassing downstream corticobulbar pathways, this approach may extend beyond brainstem or lower tract dysfunction to encompass cortical impairments such as Broca’s aphasia, thereby broadening the clinical applicability of speech BCIs.

Furthermore, the model generalized effectively to untrained syllables, demonstrating the utility of AKTs as a stable and interpretable intermediary between neural activity and phonemic categories. This feature not only enhances decoding transparency but also enables scalability beyond limited-vocabulary settings, a critical step toward open-set, real-world speech neuroprosthetics ^25,26^.

Together, these results highlight a principled path toward within-modal and cross-modal speech BCIs across speech imagery and articulation. The integration of supramodal and modality-specific articulatory representations paves the way for high-performance, low-training neural prosthetics, expanding the clinical reach of speech restoration technologies for patients across a broad spectrum of speech output disorders, particularly those with locked-in syndrome, Broca’s aphasia, or progressive conditions such as ALS ^20-22,58^.

## Methods

### Participants

A total of 9 subjects from Huashan Hospital participated in this study. All participants (median age [IQR]: 35 [32, 51] years; 4 males, 5 females; 9 left hemisphere) were eloquent brain tumor patients undergoing awake language mapping as part of their surgery. During the intraoperative language mapping, high-density electrode grids were temporarily placed onto the frontoparietal cortex to record local field potentials from the cortex, and the participants were instructed to perform the experimental tasks.

Subjects were asked to participate in the research study only if they were undergoing awake surgery with direct cortical stimulation as part of the regular clinical routine, meaning that this was deemed necessary for the safe resection of their tumor. Each participant was consented prior to the surgery, at which time it was explained in a transparent manner (as detailed in the IRB-approved written protocol/consent document) that the research task was for scientific purposes and would not directly impact their care. It was clearly articulated to each subject that participation in the research task was completely voluntary. The experimental protocol was approved by the Huashan Hospital Institutional Review Board of Fudan University (HIRB, KY2022-504). All participants gave their written informed consent prior to testing.

### Experiment paradigm

Two distinct paradigms were employed in this study. Five subjects (S1–S5) completed Paradigm 1 (**Fig. 1a**), while four subjects (S6–S9) completed Paradigm 2 (**Fig. 1b**). Both paradigms were presented using E-Prime 2.0 (Psychology Software Tools, Inc.) and were temporally synchronized with ECoG and audio recordings.

Paradigm 1 (**Fig. 1a**) consisted of two parts. In the first part, subjects were presented with the instruction "Please Press the key" and performed an ipsilateral key press task, repeated 40 times, serving as an ipsilateral hand motor baseline control. In the second part, a visual stimulus displaying a syllable (e.g., "ba") was presented on the screen for 2 seconds, followed by the instruction "Prepare to Read Aloud" for another 2 seconds. Subjects were then required to overtly produce the syllable five times (SA task) upon the appearance of a black cross symbol (go-cue) while simultaneously marking the onset of each production with a button press (with intervals of at least 1.2 s). Subsequently, the instruction "Prepare for imagery" was displayed for 2 seconds, after which subjects covertly imagined producing the syllable five times (SI task) upon the appearance of the black cross symbol (go-cue), again marking each imagined production onset with a button press (intervals > 1.2 s). The ipsilateral key press served as the reference time point for aligning neural, audio, and feature matrix data.

Paradigm 2 (**Fig. 1b**) introduced an auditory cueing component. Subjects first listened to a syllable auditory stimulus (e.g., "ga") repeated five times. The syllable auditory stimuli were extracted from recordings of the subjects’ own voices, obtained during preoperative training. This was followed by the instruction "Prepare to Read Aloud", accompanied by a visual display of the syllable (2 s). Next, a gray cross symbol and a black cross symbol alternated every 1 s, and subjects were instructed to overtly produce the syllable each time the black cross appeared, repeating this process five times (SA task). The syllable imagery task followed the same procedure: after the instruction "Prepare for imagery" and the visual presentation of the syllable (2 s), subjects covertly imagined producing the syllable in sync with the black cross symbol, repeating the task five times (SI task). The appearance of the black cross (go-cue) served as the reference time point for subsequent neural, audio, and feature matrix segmentation.

Regarding syllable selection, Paradigm 1 included six syllables: /ba/, /da/, /ga/, /bu/, /du/, and /gu/, each repeated 60 times in both the SA and SI tasks. Paradigm 2 included eight syllables: /ba/, /da/, /ga/, /pa/, /ta/, /ka/, /sa/, and /sha/, each repeated 45 times in both the SA and SI tasks.

Four methodological controls were implemented to eliminate potential articulatory muscle activation during the speech imagery task: (1) During preoperative training, subjects were required to complete a full iteration of the intraoperative paradigm and were explicitly instructed to avoid articulatory movements or voicing during SI tasks. (2) During preoperative training, a researcher (Z.Z.) closely monitored the subjects, providing immediate corrections to ensure the absence of visible articulatory movements and audible speech during the SI task. (3) During the intraoperative SI task, real-time video recordings of subjects’ orofacial movements and corresponding audio recordings were monitored and acquired using the Brain Mapping Interactive Stimulation System (Shenzhen Sinorad Medical Electronics Co. Ltd) ^77^ to confirm the absence of motor and auditory output. (4) For the last four subjects (S6–S9), intraoperative electromyographic (EMG) recordings were obtained using needle electrodes placed on orofacial muscles (e.g., orbicularis oris, mylohyoid) to ensure that the included SI trials exhibited no detectable EMG activity exceeding the predefined threshold indicative of articulatory muscle engagement.

### ECoG data acquisition and preprocessing

During the experimental tasks, neural signals were recorded from one or two 128-channel ECoG grids (8 × 16, 3 or 4 mm spacing) using a multichannel amplifier optically connected to a digital signal processor (Tucker-Davis Technologies). The local field potential at each electrode contact was amplified and sampled at 3052 Hz.

The raw voltage waveforms were visually inspected, and channels with undetectable signal variation relative to noise or exhibiting continuous epileptiform activity were excluded. Time segments containing electrical or movement-related artifacts were manually identified and removed. The remaining signals were then notch-filtered to eliminate line noise at 50 Hz, 100 Hz, and 150 Hz.

To extract frequency-specific neural activity, band-pass filtering was applied to isolate the following frequency bands: theta (4–8 Hz), alpha (8–12 Hz), beta1 (12–24 Hz), beta2 (24–40 Hz), low gamma (40–70 Hz), and high gamma (70–150 Hz) ^21,33^. The analytic amplitude (envelope) of each band was computed using the Hilbert transform and smoothed using a Gaussian filter. The signal for each frequency band was obtained by averaging the analytic amplitude across eight logarithmically spaced sub-bands within the specified frequency range^33^. Finally, the signal was down-sampled to 100 Hz and z-scored using the entire recording block for normalization ^31^.

### Cortical surface extraction and electrode localization

We used FreeSurfer to generate pial surface reconstructions from preoperative T1-weighted MRI scans. To localize electrode positions, the three-dimensional coordinates of the grid corners were recorded intraoperatively using the Medtronic neuronavigation system. These corner electrodes were then co-registered to the preoperative MRI, with intraoperative photographs serving as additional reference points. The remaining electrode positions were subsequently estimated using the “img_pipe” package in Python, which interpolates and extrapolates electrode locations based on the recorded corner coordinates ^78^. Electrode positions in individual spaces were normalized to the MNI 152 (2009, asymmetric) template, and anatomical labels were assigned based on the Desikan-Killiany ^79^ and Human Connectome Project’s multi-modal parcellation (MMP) atlases ^36^.

### Responsive electrode selection

After the neural activity computation in each frequency band, response electrodes for both SA and SI were identified. The average time difference between syllable onset in the audio (SA task) and the reference time point (ipsilateral key press in Paradigm 1 or the go-cue appearance in Paradigm 2) was calculated. This value was then used to align the ECoG data to the syllable onset in the SA modality or internal syllable onset in the SI modality (set as 0 ms), followed by segmentation and averaging across tasks.

The neural activity between -500 ms and -400 ms served as baseline resting-state activity. A two-sample *t*-test (Bonferroni correction, *α* = 0.05/160, where 160 is the number of time points tested) compared average neural activity between baseline and each time point from - 400 ms to +1200 ms. Electrode responses were considered significant if at least 100 ms (10 consecutive time points) showed significant differences. To control for motor responses in Paradigm 1, electrodes responsive to the ipsilateral key press task (control) were excluded from further analysis.

### Speech feature extraction

We employed a similar Praat parameter extraction approach in line with our previous research ^80,81^. We extracted the pitch contour (log F_0_, F_0_ = fundamental frequency) of each syllable from the audio recorded during the SA modality with an autocorrelation method in Praat (https://www.fon.hum.uva.nl/praat/, Version 6.1.01) ^82^, which was subsequently used to compute the one-dimensional trajectory for the larynx. Additionally, we addressed halving and doubling errors during the extraction process. Individual pitch minimum and maximum values were determined for each participant, and a timestep of 0.01s was employed. All other parameters adhered to Praat’s default settings.

The onset (or internal onset for SI modality) of phonemes for all syllables was manually labeled in Praat by the researchers (Z.Z., Z.W., and Y.L.) and subsequently categorized into four places of articulation (POA) classes: (1) Labial: /b/, /p/; (2) Coronal: /d/, /t/, /s/, /sh/; (3) Dorsal: /g/, /k/; (4) Vocalic: /a/, /u/.

### Speaker-Independent Acoustic-to-Articulatory Inversion (AAI)

Following prior studies ^28,83^, we applied a Speaker-Independent Acoustic-to-Articulatory Inversion (AAI) algorithm to convert synchronized audio recorded during the SA modality into 13-dimensional articulatory kinematic trajectories (AKTs) for each syllable. Previous studies have demonstrated a strong positive correlation between these articulatory kinematic trajectories and electromagnetic midsagittal articulography recordings ^28^. These AKTs captured two-dimensional trajectories (x and y directions) for six articulators—upper lip (UL), lower lip (LL), lower incisor/jaw (LI), tongue tip (TT), tongue body (TB), and tongue dorsum (TD)— and a one-dimensional trajectory for the larynx (L), represented by the scaled log F_0_ (fundamental frequency) and normalized to a range of -1 to 1 a.u. For SI tokens, we used the corresponding SA tokens’ AKTs from the same trial as representations of internal AKTs. For detailed code of the Speaker-Independent AAI algorithm, please refer to: github.com/articulatory/articulatory.

### Encoding model

We used time-delayed linear encoding models, known as temporal receptive field models ^84^, to evaluate what features drive the neural activity in frontoparietal regions during SA and SI modalities. Temporal receptive field (TRF) models predict neural activity using speech-related features (AKTs or POAs) in a time window (-400 ms to + 400 ms) around the neural activity. In particular, we fit the linear model for each electrode:

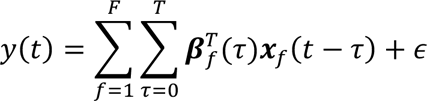

Parameter y is the neural activity recorded from the electrode, ***x***_*f*_(*t* − *τ*) is the stimulus representation vector of feature set *f* at time *t* − *τ*, ***β***_*f*_(*τ*) is the regression weights for feature set f at time lag *τ*, and *ϵ* is the Gaussian noise.

To prevent overfitting, we employed L2 regularization and cross-validation. The data were divided into three mutually exclusive sets, comprising 80%, 10%, and 10% of the samples. The 80% set was used for training, the 10% set for optimizing the L2 regularization hyperparameter, and the remaining 10% set for testing. Model performance was evaluated by the coefficient of determination (*R*^2^) between the actual and predicted neural activity values on the held-out test data. This procedure was repeated using five-fold cross-validation, with the final model performance reported as the mean *R*^2^ across all folds, reflecting the total variance explained by the model.

In the full TRF model for AKTs, the feature matrix of the 13-dimensional trajectories for seven articulators was included. To compare the performance of the AKT models across all frequency bands in both the SA and SI modalities, we first conducted a Kruskal-Wallis *H*-test with a significance threshold of *p* < 0.05. Subsequently, pairwise Mann-Whitney *U* tests were performed, with a significance threshold of *p* < 0.05 after the Benjamini-Hochberg correction. To quantify the unique contribution of each articulator, we fitted TRF models excluding each articulator in turn and computed the change in *R*^2^ (Δ*R*^2^) between the full and reduced models.

We also trained a full TRF model incorporating POA cluster-based features, where the presence or absence of each phoneme’s POA feature at its onset was marked using binary indicators (1 for presence, 0 for absence). The unique contribution of each POA was also quantified as the Δ*R*^2^ between the full and reduced models.

Building upon the electrodes exhibiting significant activity in the beta1 band, we further excluded electrodes that did not encode AKTs. Specifically, we removed electrodes with an *R*^2^ value of ≤ 0.01 in both the SA- and SI-AKT modalities, corresponding to the 50% threshold of the *R*^2^ distribution across all responsive electrodes (**Extended Data Fig. 4b**).

### K-means clustering for encoding electrode classification

For dual-responsive electrodes, we aimed to determine whether beta1 neural activity encodes AKTs similarly across SA and SI modalities. To address this, we first calculated the *R*^2^difference between the SA-AKT and SI-AKT models for each dual-responsive electrode. We applied the Duda-Hart test to evaluate whether the differences surpassed the threshold (Duda-Hart statistic *d* > 1.645, *p* < 0.05), which would indicate the necessity for classification into at least two clusters ^85^. Following this, we performed unsupervised K-means clustering (KMeans from scikit-learn python package), iterating the number of clusters from 2 to 10, and selected the optimal number of clusters based on the highest silhouette score ^86^. For each cluster, we analyzed the similarity in performance (*R*^2^ values) between the SA-AKT and SI-AKT models for the electrodes within the cluster using Pearson correlation analysis. Differences in performance were assessed using the Wilcoxon signed-rank test (two-sided). A *p*-value of < 0.05 was considered statistically significant for both analyses.

### Cosine similarity of encoding patterns across SA and SI modalities

To assess whether electrodes within specific clusters exhibit shared AKT encoding patterns, we constructed encoding matrices for each modality. Each matrix entry represented each articulator’s unique contribution (Δ*R*^2^) across electrodes, scaled to a range of 0 to 1 arbitrary unit. We then calculated the cosine similarity between the encoding matrices for the two modalities. A permutation test was performed to evaluate statistical significance: electrode labels within each encoding matrix were randomly shuffled, and the cosine similarity was recalculated for 10,000 iterations to generate a null distribution. The observed cosine similarity was compared to this distribution, with significance determined if the *p*-value was < 0.05. The same approach was applied to assess the cosine similarity of POA encoding patterns between the SA and SI modalities.

### Hierarchical clustering

Agglomerative hierarchical clustering was performed using Ward’s method. Clustering was applied along the electrode dimension for AKT (**Fig. 2s, 2u**) and POA encoding matrices (**Extended Data Fig. 5a, 5c**).

To examine clustering patterns of AKTs across phonemes, we constructed a phoneme-specific AKT displacement matrix (**Fig. 4a**). For each phoneme, we calculated the average AKTs across all subjects, defining the direction of maximum deviation from the resting state as the primary movement direction (positive). The maximum displacement of each phoneme’s 13 AKTs was then computed, with a displacement sign assigned based on alignment with the primary movement direction. The resulting AKT displacement matrix for each phoneme was normalized to a range of 0 to 1 arbitrary unit. Hierarchical clustering was performed on both the phoneme and AKT dimensions using SciPy and Seaborn Python packages ^87^.

### Spatial distribution of encoding electrodes

To investigate the spatial distribution of encoding electrodes across different categories (SA-SI shared, SA-specific, and SI-specific), kernel density estimation (KDE) was performed in the y-z plane of MNI space. KDE was applied with Scott’s bandwidth selection method to construct probability density distributions for each encoding category. The distributions were thresholded at 0.5 cumulative density (a.u.).

The KDE maps were processed using a maximum filter from the SciPy package (scipy.ndimage.maximum_filter) to identify local density peaks. Statistical comparisons of electrode distributions in MNI space were performed using the two-sided Kolmogorov-Smirnov test to assess differences along the y-axis and z-axis (significant if *p* < 0.05).

Similar KDE maps were generated for each POA across different encoding categories, with electrode positions weighted by their unique contribution (Δ*R*^2^) to the full POA model. To evaluate differences in weighted KDE distributions between POAs along the y-axis and z-axis, we applied a permutation test based on the Wasserstein distance. For each permutation (5000 iterations), electrode positions and weights (Δ*R*^2^) were shuffled, and the Wasserstein distance was recalculated between POA distributions. The *p*-values were derived from the permutation distribution and corrected for multiple comparisons using the Benjamini-Hochberg procedure, with significance at *p* < 0.05 after correction.

### Temporal distribution of encoding electrodes

To investigate the temporal distribution of encoding electrodes, we calculated the onset and peak times during syllable production and imagery tasks. For each time point within the -400 ms to +1200 ms window, an independent two-sample t-test was performed comparing task-related beta1 activity to baseline. The onset time was defined as the first time point with a sustained *p*-value < 0.01 for at least 100 ms. The peak time was determined as the point of maximal beta1 amplitude following onset. Differences in onset and peak times across encoding categories were evaluated using the Mann-Whitney *U* test, with significance at *p* < 0.05.

### Triple-stream parallel neural decoding framework

To determine whether beta1 activity across discrete multi-electrode sites could be leveraged for speech decoding, particularly for inner speech decoding in the SI modality, we developed a triple-stream parallel neural decoding framework (**Fig. 5a**) using the Pytorch and Keras packages ^88,89^. This framework decodes beta1 activity from SA- and SI-responsive electrodes into three highly interrelated components: syllable classification (syllable classifier, **Fig. 5b**), audio synthesis (speech synthesizer, **Fig. 5c**), and AKT reconstruction (articulatory movement synthesizer, **Fig. 5d**). Notably, since the SI modality does not generate auditory or motor output, we incorporated the speech feature matrix from the corresponding SA trial as an internal representation of speech to train the decoding model. For all three streams, input data were structured as channels × time steps, where channels corresponded to SA- or SI-responsive electrodes, depending on the task modality. The time window extended from 0.5 s before to 1.5 s after syllable onset, comprising 200 time steps at a sampling rate of 100 Hz. For the articulatory movement synthesizer, the first and last four time steps were excluded to mitigate the edge effects of AKTs.

**The syllable classifier (Fig. 5b)** is an adapted 34-layer Residual Network (ResNet-34) architecture ^90^. The first convolutional layer employed a 7 × 7 kernel, followed by a batch normalization layer and a 2D max-pooling layer. Four residual blocks (ResBlocks) were subsequently applied. All convolutional layers in the decoder utilized Rectified Linear Units (ReLU) as activation functions ^91^. A 40% dropout layer was included to mitigate overfitting. The network was trained to minimize joint cross-entropy loss using the stochastic gradient descent (SGD) optimizer, with an initial learning rate of 0.01, a decay rate of 1×10⁻10^6^, the momentum of 0.9, and Nesterov acceleration enabled. Optimization was halted when validation loss ceased decreasing. The classifier’s performance was evaluated using syllable decoding accuracy, employing a 10-fold cross-validation scheme to determine the average performance across folds.

**The speech synthesizer (Fig. 5c)** employs a deep convolutional neural network architecture composed of four sequential modules, each consisting of a Conv2D layer, a ResBlock ^90^, and a 2D max-pooling layer, followed by a fully connected layer. All convolutional kernels were set to 3 × 3. To assess the quality of synthesized speech waveforms, we used Mel-cepstral distortion (MCD) as an objective measure. MCD quantifies the error in Mel-Frequency Cepstral Coefficients (MFCCs) ^30,92^ and was calculated as follows:

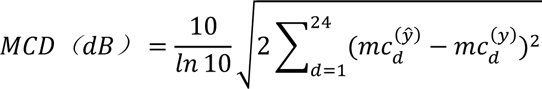

where *d* represents each MFCC dimension (0 < *d* < 25), *ŷ* is the synthesized speech, and *y* is the actual participant-produced acoustic signal ^92^. Performance evaluation used a 10-fold cross-validation strategy to determine the average MCD across folds.

**The articulatory movement (Fig. 5d)** synthesizer employs a vector-quantized variational autoencoder (VQ-VAE) ^93^. In the first stage, an encoder block composed of residual and non-local blocks extracts features from the ECoG signals. In the second stage, extracted features are mapped onto a 256-dimensional articulatory codebook, yielding a discretized codebook representation. In the third stage, the codebook mapping is fed into a decoder block (also consisting of residual and non-local blocks) ^90^ to reconstruct articulatory movements with dimensions of 13 × 192. The network was optimized to minimize mean squared error loss using the Adam optimizer, with an initial learning rate of 2.25×10^-5^, *β₁* = 0.5, *β₂* = 0.9, and *ε* = 1×10^-8^. Performance evaluation was conducted using a 5-fold cross-validation strategy. The dataset was randomly divided into ten subsets, with eight used for training, one for validation, and one for testing. For each fold, performance was assessed based on the mean Pearson’s correlation coefficient (*r*) across 13-dimensional AKTs. The final performance was reported as the average across all five folds.

To validate the robustness of this triple-stream parallel decoding framework, we conducted tests on both SA and SI modalities in nine participants. Median syllable decoding accuracy was compared against chance levels (1/6 for six-class tasks in subjects S1-S5; 1/8 for eight-class tasks in subjects S6-S9). Median MCD was compared against the 8 dB threshold, the upper limit considered acceptable for voice recognition systems ^30^. Additionally, median *r* values were compared against chance levels, defined as the correlation between AKTs synthesized by an untrained, randomly initialized model and actual AKTs. All statistical comparisons were conducted using the Mann-Whitney *U* test. Modality differences were assessed using the Wilcoxon signed-rank test, with *p* < 0.05 considered statistically significant.

To further investigate whether cross-modal articulatory movement decoding could be achieved using SA-SI shared electrodes, we trained four articulatory movement synthesizers: SA2SA, SA2SI, SI2SI, and SI2SA (naming format indicates "training set to testing set," e.g., SA2SI denotes training with SA modality beta1 activity and testing with SI modality beta1 activity to evaluate the correlation between synthesized and ground-truth movements). Differences among these four models were assessed using the Kruskal-Wallis *H* test, with *p* < 0.05 considered statistically significant.

Lastly, we examined whether the articulatory movement synthesizer effectively captured a stable relationship between beta1 activity and AKTs, allowing for extrapolative generalization. Specifically, the model’s ability to predict movement trajectories of syllables excluded from the training set was assessed using a Leave-One-Syllable-Out strategy. In each iteration, one syllable was excluded from training, and the model was trained on the remaining syllables. The excluded syllable was then used as a testing set, and the correlation (*r*) between predicted and actual AKTs was computed. Chance-level *r* values were determined using an untrained, randomly initialized model. Median *r* values and modality differences were statistically assessed using the Mann-Whitney *U* and Wilcoxon signed-rank tests, respectively.

## Data availability

The data set generated during the current study will be made available from the authors upon reasonable request.

## Code availability

The completely developed code that operates on the full data set will be made available from the corresponding authors upon reasonable request.

## Acknowledgments

Dr. Junfeng Lu is supported by STI 2030—Major Projects (2022ZD0212300) and National Natural Science Foundation of China (32371146). Dr. Zehao Zhao is supported by the Postdoctoral Fellowship Program of CPSF (GZB20240661). Dr. Jinsong Wu is supported by Innovation Program of Shanghai Municipal Education Commission (2023ZKZD13). Dr. Yuanning Li is supported by the National Natural Science Foundation of China (32371154), Shanghai Rising-Star Program (24QA2705500), Shanghai Pujiang Program (22PJ1410500, Y.L.), and the Lingang Laboratory (LG-GG-202402-06). The computations in this research are supported by the HPC Platform of ShanghaiTech University.

## Author contributions

Z.Z., Z.W., Y.Liu., Y.Li., J.L., and J.W. conceived and designed the study. Z.Z., Z.W., Y.Li., and Y.Y. constructed the temporal receptive field model and performed statistical analyses. Z.Z., Z.W., Y.Li., and Y.Liu. developed and optimized the decoding framework. Z.Z., Y.Liu., Y.Q., J.L., and J.W. collected neural and behavioral data. J.L. and J.W. coordinated session logistics, managed equipment, and performed cortical electrode placement. X.G., B.Y., S.X.T., and X.T. provided guidance on neurolinguistic experimental design and contributed to manuscript revision. J.W. supervised the overall project and provided strategic guidance. The study was jointly supervised by G.C., Y.Li., J.L., and J.W. Z.Z. wrote the initial draft with input from Y.Liu. and Z.W. All authors reviewed and edited the manuscript.

