## Extended Data Figs. 1-8 for "Neural hierarchy for coding articulatory dynamics in speech imagery and production"

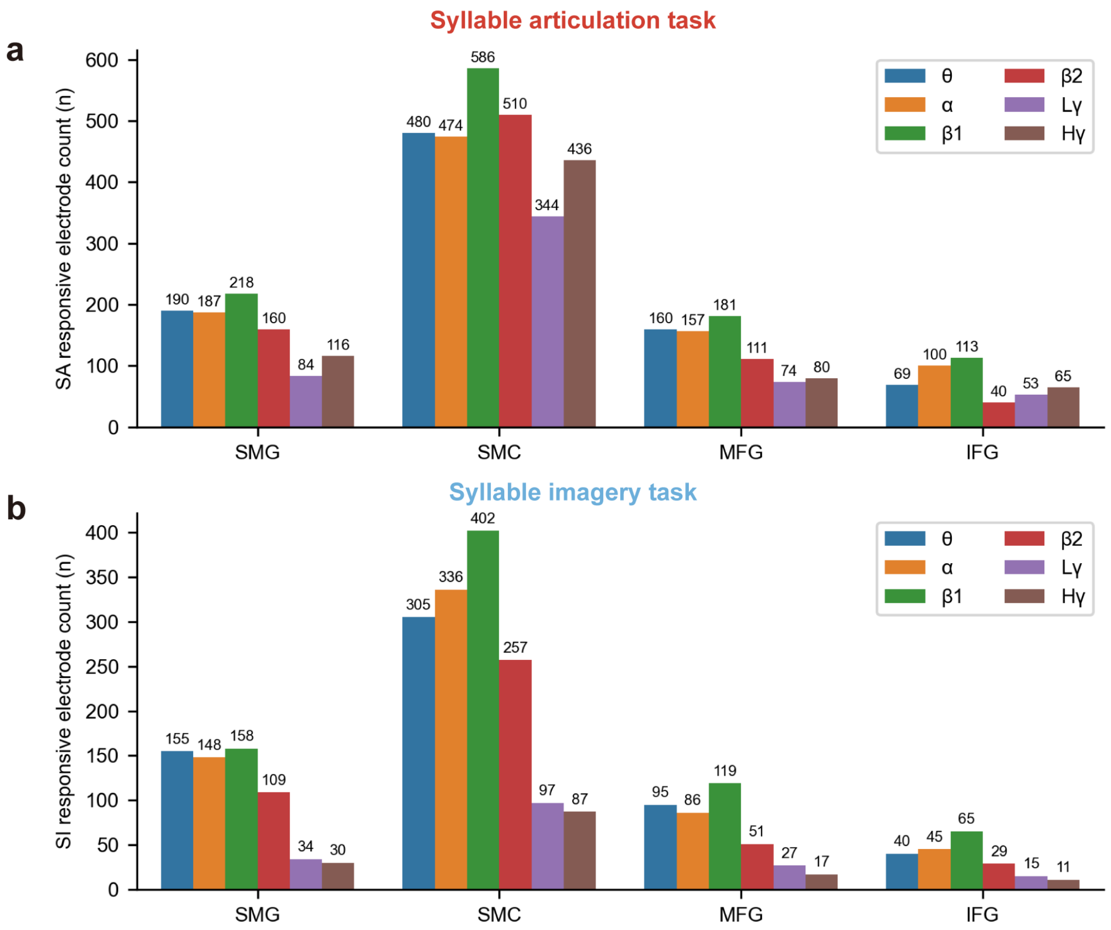


**Extended Data Fig. 1 | Number of responsive electrodes across frequency bands within different brain regions during syllable articulation (SA) and syllable imagery (SI).**

**a-b,** Bar plots showing the distribution of responsive electrodes across frequency bands in different brain regions for SA **(a)** and SI **(b)**: theta (θ, 4-8 Hz), alpha (α, 8-12 Hz), beta1 (β1, 12-24 Hz), beta2 (β2, 24-40 Hz), low gamma (Lγ, 40-70 Hz), and high gamma (Hγ, 70-150 Hz). SMG = supramarginal gyrus, SMC = sensorimotor cortex, MFG = middle frontal gyrus, IFG = inferior frontal gyrus.

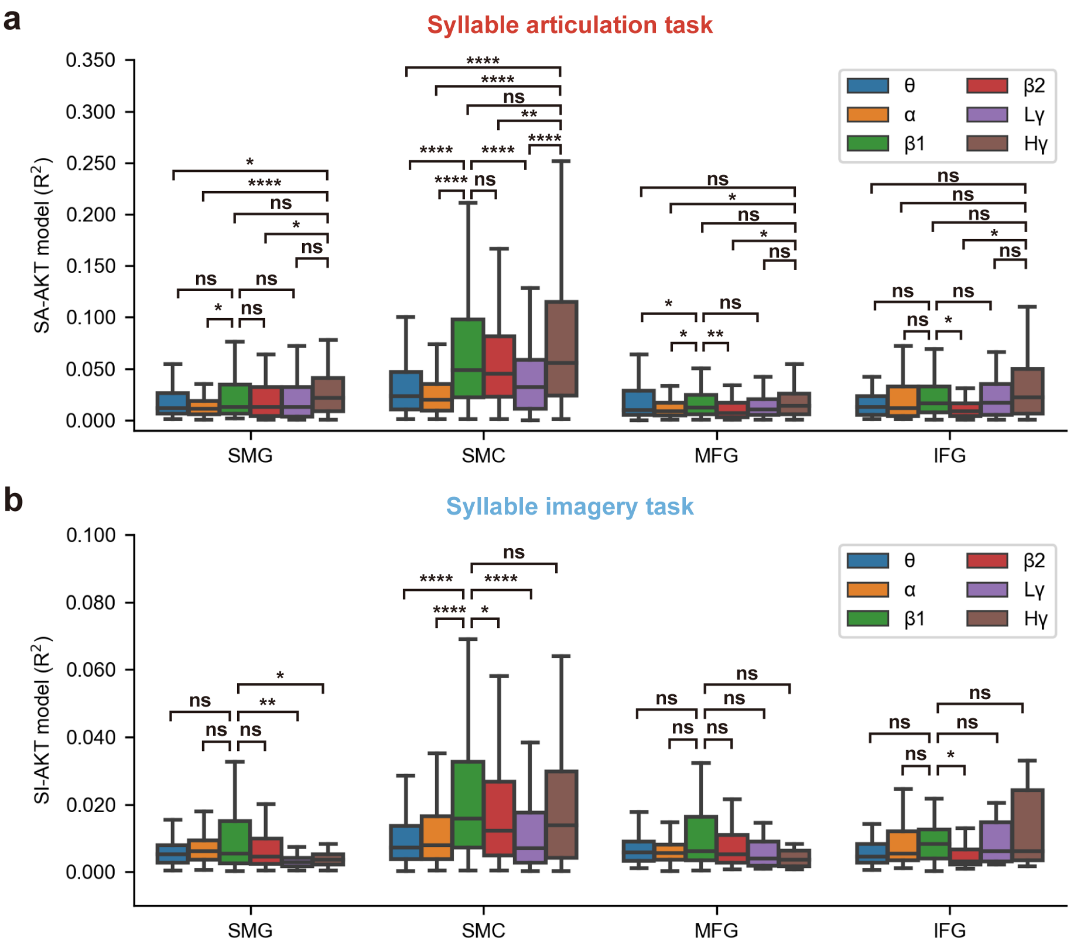


**Extended Data Fig. 2 | Frequency band preference for articulatory kinematic trajectory (AKT) encoding within different brain regions during SA and SI tasks.**

**a-b,** Box plots depicting AKT model performance ($R^2$) across frequency bands within distinct brain regions for SA **(a)** and SI **(b)**: theta (θ, 4-8 Hz), alpha (α, 8-12 Hz), beta1 (β1, 12-24 Hz), beta2 (β2, 24-40 Hz), low gamma (Lγ, 40-70 Hz), and high gamma (Hγ, 70-150 Hz). *$p$ < 0.05, **$p$ < 0.01, ****$p$ < 0.0001, ns: non-significant (two-sided Mann-Whitney $U$ tests with Benjamini-Hochberg correction). Comparisons: SA (Hγ or β1 vs. other bands) and SI (β1 vs. other bands).

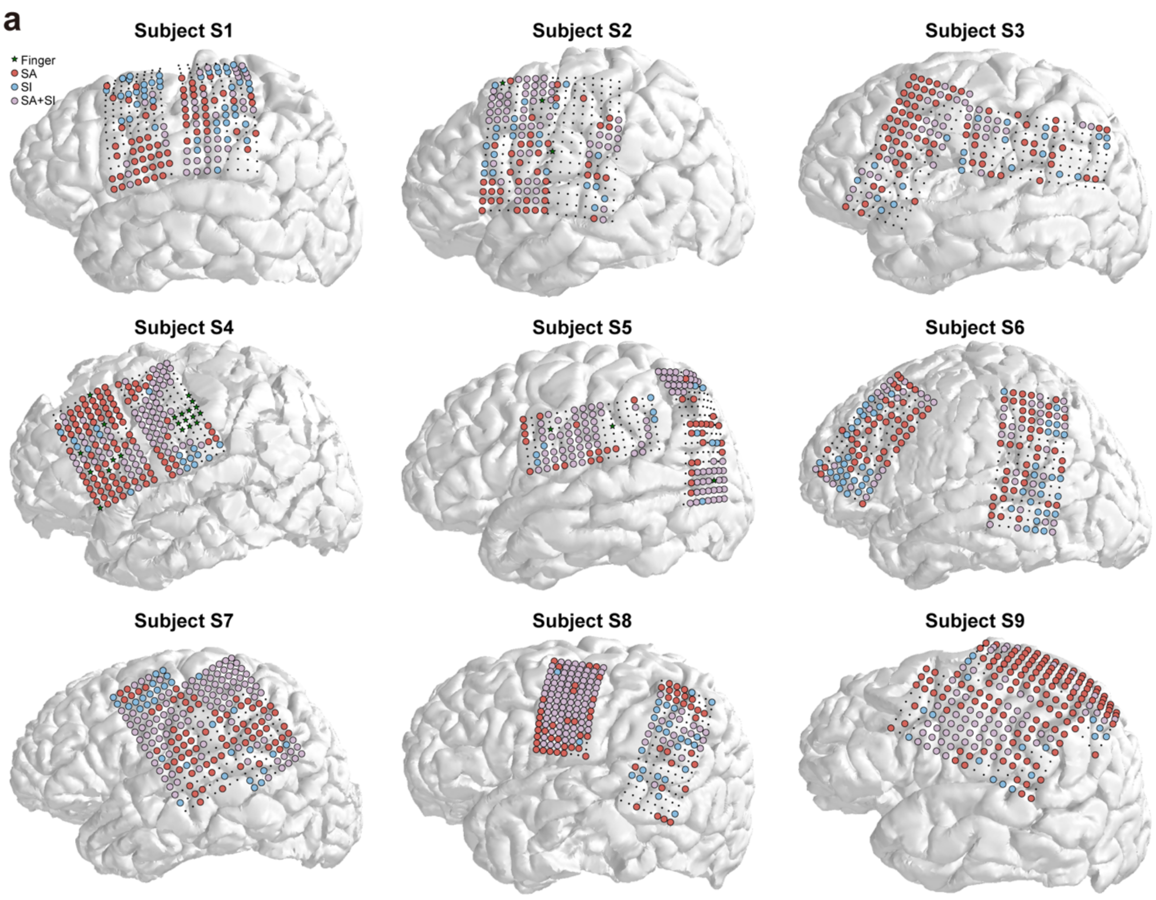


**Extended Data Fig. 3 | Individual-level spatial distribution of shared and modality-specific responsive electrodes in SA and SI, mediated by frontal-parietal beta1 activity.**

**a,** Individual brain reconstructions for subjects S1–S9, illustrating electrode coverage. Green stars: ipsilateral hand motor-responsive electrodes; light red, light blue, and light purple circles: SA-only, SI-only, and SA-SI dual-responsive electrodes, respectively.

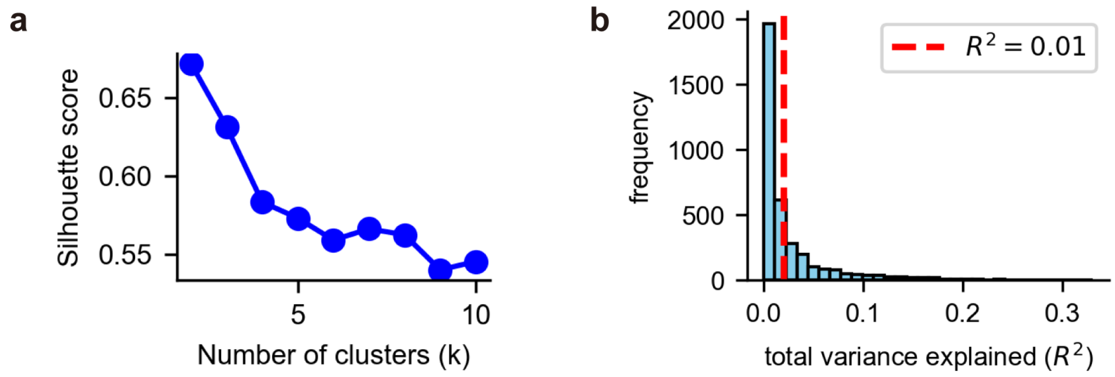


**Extended Data Fig. 4 | Silhouette scores from K-means clustering analysis and distribution of AKT encoding performance.**

**a,** Silhouette scores for varying numbers of clusters (*k* = 2–10), with the maximum score observed at *k* = 2. **b,** Histograms of AKT encoding performance ($R^2$) for all responsive electrodes. The red dashed line indicates $R^2$ = 0.01, corresponding to the 50% threshold.

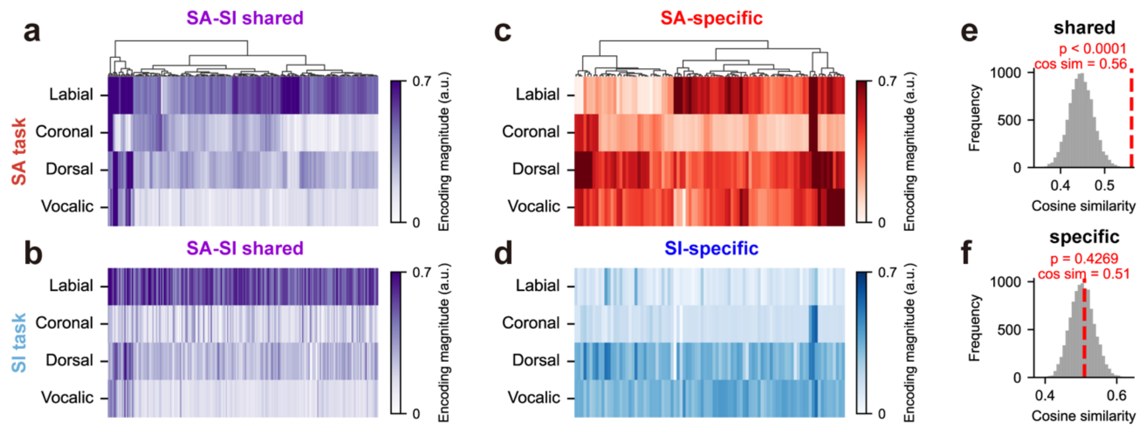


**Extended Data Fig. 5 | Shared and specific place of articulation (POA) encoding in dual-responsive electrodes across SA and SI modalities.**

**a, b, e,** POA-encoding patterns, computed by the TRF model for each SA-SI shared encoding electrode during SA (**a**) and SI (**b**) tasks. Electrodes in **(a)** are organized by hierarchical clustering; the same order is maintained in **(b)**. Color intensity represents the unique contribution ($\Delta R^2$) to the full POA model, scaled to a range of 0 to 1 a.u. **e,** Histogram from permutation testing shows significant similarity in POA-encoding patterns between SA-SI shared encoding electrodes during SA and SI tasks (red dashed line, *cosine similarity* = 0.561, *p* < 0.0001). **c, d, f,** For overlapping electrodes categorized as SA-specific (**c**, red, during syllable articulation) and SI-specific (**d**, blue, during syllable imagery), the histogram from permutation testing (**f**) shows no significant similarity in POA-encoding patterns (red dashed line, *cosine similarity* = 0.510, *p* = 0.4239). Electrodes in **(c)** are organized by hierarchical clustering; the same order is used in **(d)**. Color intensity represents the unique contribution ($\Delta R^2$) to the full POA model, scaled to a range of 0 to 1 a.u.

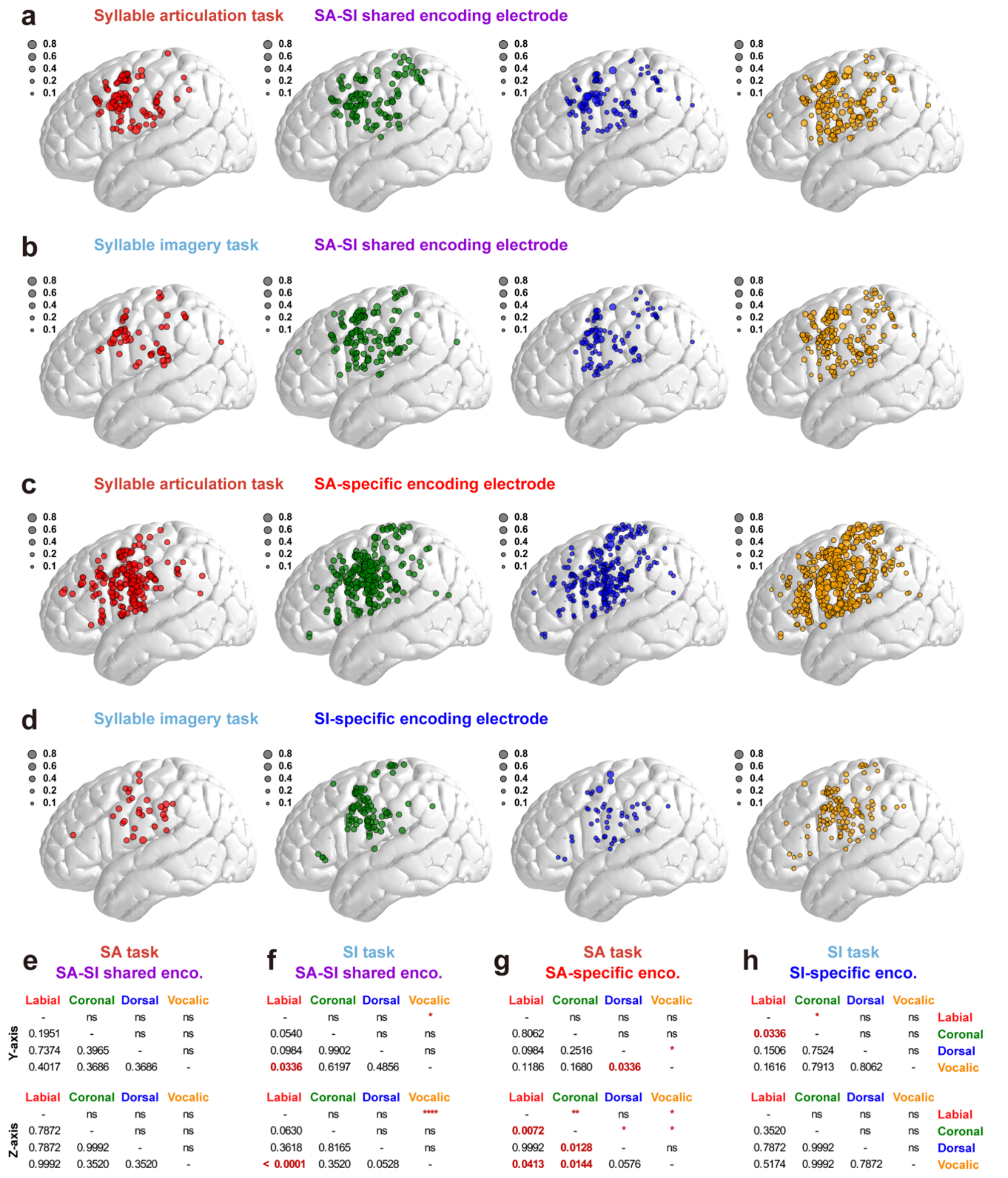


**Extended Data Fig. 6 | Scatter plots of the unique contributions of POAs in SA-specific, SI-specific, and SA-SI shared encoding electrodes during SA and SI modalities.**

**a-d,** Weighted scatter plots for the four POA categories during the syllable articulation task (**a, c**) and the syllable imagery task (**b, d**), visualized for SA-SI shared (**a, b**), SA-specific (**c**), and SI-specific (**d**) encoding electrodes. The size of the scatter points is weighted by the $\Delta R^2$ values (normalized to a scale of 0–1) of the POAs. Each color represents a distinct POA: red for labial, green for coronal, blue for dorsal, and yellow for vocalic. **e-h,** The tables display the results of pairwise permutation tests comparing the weighted probability density distributions of POAs along the Y and Z axes of MNI space. The lower left triangle contains Benjamini-Hochberg corrected *p*-values, while the upper right triangle shows the corresponding statistical significance symbols (ns: not significant, $*p < 0.05$, $**p < 0.01$, $****p < 0.0001$).

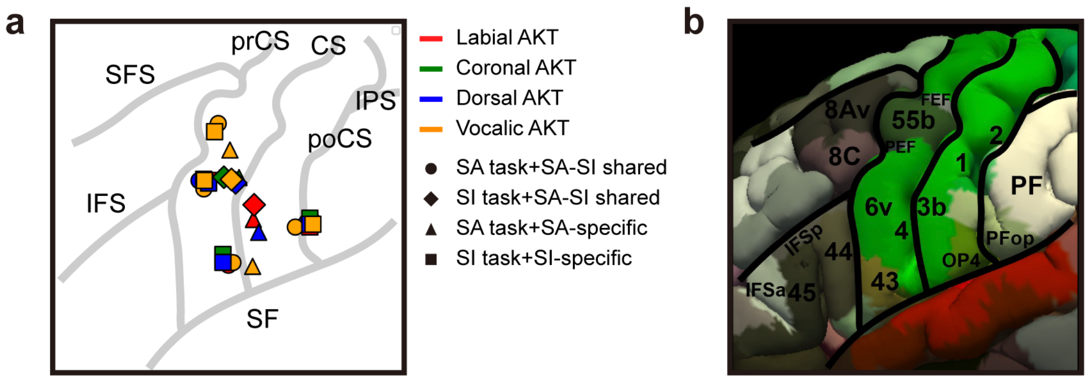


**Extended Data Fig. 7 | Peak locations of POA encoding across SA-specific, SI-specific, and SA-SI shared electrodes during SA and SI modalities.**

**a,** Peak points of weighted probability density distributions of POAs along the Y and Z axes of MNI space. Different colors represent distinct POAs, while different symbols denote the classification of encoding electrodes, as indicated in the figure. **b,** Subdivisions of the frontal and parietal lobes based on the multi-modal parcellation (MMP) atlas from the Human Connectome Project.

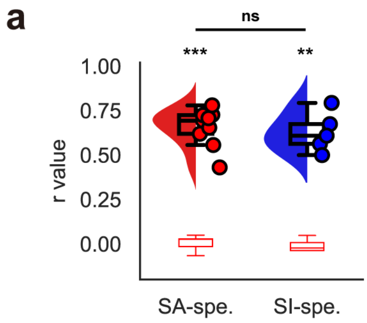


**Extended Data Fig. 8 | AKT decoding performance across SA-specific and SI-specific electrodes.**

**a,** Violin plots showing decoding performance of synthesized AKTs, quantified by the mean Pearson's *r* between predicted and ground-truth AKTs, for SA-specific ($N$ = 9 subjects) and SI-specific ($N$ = 5 subjects) electrodes. In the remaining four subjects, decoding could not be performed due to insufficient SI-specific electrodes for model training. The red box indicates the chance level. **$p < 0.01$, ***$p < 0.001$, Mann–Whitney *U* test against chance; ns, not significant for comparison between SA- and SI-specific electrodes (Mann–Whitney *U* test). Spe., specific.
